# Recognition-dependent activation of the RRS1-R/RPS4 immune receptor complex

**DOI:** 10.1101/2025.04.11.646618

**Authors:** Hee-Kyung Ahn, Guanghao Guo, Jan Skłenar, Shania Pin Yin Keh, Sung Un Huh, Michelle T. Hulin, Hayden Burdett, Maria Sindalovskaya, Jihyeon Choi, Nitika Mukhi, He Zhao, Lena Knorr, Mark J. Banfield, Frank L.H. Menke, Wenbo Ma, Jonathan D.G. Jones

## Abstract

The Arabidopsis TIR-NLR immune receptors RPS4 and RRS1 function together to enable recognition of multiple effector proteins including AvrRps4 and PopP2. We show here that both in the presence and absence of effector, RPS4 and RRS1 form an oligomer that does not change in size upon effector provision. Oligomer formation involves interactions between the RPS4 and RRS1 TIR domains and requires nucleotide binding capacity in RPS4. RPS4 mutants that lose TIR domain NADase activity abrogate immune activation but retain oligomerization. A cysteine residue in the RPS4 LRR domain contributes to oligomer stabilization. We propose that upon effector recognition, conformational changes in the complex relieve inhibition of RPS4 TIR domains by RRS1 TIR domains, enabling proximity between RPS4 TIR domains to create NADase activity.

## Introduction

Plant diseases significantly reduce crop yields and plant breeders deploy *Resistance (R)* genes that reduce such losses. Many *R* genes are now identified, and most encode intracellular Nucleotide-binding, Leucine-rich repeat [LRR] immune Receptors (NLRs)^1^ that directly or indirectly detect pathogen effectors secreted into plant cells. NLR genes are found in plants and animals^2–4^, and upon ligand perception, NLRs oligomerize into complexes, mediated mainly by the canonical NB-ARC (Nucleotide-Binding domain in APAF1, Resistance, CED-4) domain or NACHT domain (for animal NLRs)^5^ and activate N-terminal signalling domains by imposing induced proximity^6^.

Plant NLRs carry Coiled-Coil (CC), CC-RPW8 (CC_R_), or TIR (Toll, Interleukin-1 Receptor, Resistance) N-terminal signalling domains, and their induced proximity results in various outcomes. Activation of CC domain (CC-NLRs) or CC_R_-carrying NLRs (CC_R_-NLRs) results in cation channel formation, Ca^2+^ influx and a localized cell death known as HR (Hypersensitive Response)^7,8^. Structural analysis of CC-NLR ZAR1^9^ and Sr35^10^ receptors revealed pentameric protein complexes formed upon effector recognition. In contrast, activation of the CC-NLRs NRC4 and NRC2 results in hexamer formation^11,12^. Refined bioinformatic and phylogenomic analyses have revealed previously unappreciated diversity within the CC-NLR class^13^.

Structural analysis of activated TIR-NLRs Roq1 (Recognition of XopQ 1) and RPP1 (Recognition of *Peronospora parasitica* 1) revealed four TIR-NLR protomers bound to four protomers of their cognate effectors, XopQ (*Xanthomonas* Outer Protein Q) or ATR1 (*Arabidopsis Thaliana* Recognized 1), respectively^14,15^. In these structures resolved by cryo-EM, TIR domains are assembled into a ‘dimer of dimers’ configuration, revealing two NADase active sites from each TIR-domain dimer. Mammalian and bacterial TIR domains can also show NADase activity^16^. In all cases, the catalytic glutamate residue and an aspartate residue in the region known as the BB-loop are crucial for NADase activity^17–19^.

NADase activation results in depletion of NAD+ and cytotoxicity in mammals and bacteria^16,20^. In plants, TIR domain NADase activity catalyses NAD+ conversion into small molecules that are recognized by the EDS1 (Enhanced Disease Susceptibility 1)/PAD4 (Phytoalexin Deficient 4) or EDS1/SAG101 (Senescence Associated Gene 101) heterodimers^21,22^. Small molecule-bound EDS1/PAD4 or EDS1/SAG101 heterodimers then interact with CC_R_-NLRs ADR1 (Activated Disease Resistance 1) or NRG1 (N-Requirement Gene 1)^23,24^. The EDS1/SAG101/NRG1 oligomer plays a key role in HR whereas EDS1/PAD4/ADR1 promotes bacterial resistance in Arabidopsis, and both HR and resistance in Solanaceae plants such as *Nicotiana benthamiana*^25^.

Some NLRs require other NLRs for function^26^. The CC_R_-NLRs and the NRC (NLR required for cell death) clade of NLRs provide a ‘helper’ function required for signalling upon recognition by ‘sensor’ NLRs. Effector recognition by the ‘sensor’ NLRs activates ‘helper’ NLRs without formation of a stable association between them^27,28^. The mammalian NLR NLRC4 forms a heterooligomer upon activation by sensor NLR NAIP proteins^29^ that bind ligand and then interact with and drive oligomerization of NLRC4. Paired NLRs are different from NLRC4/NAIPs and from most plant sensor/helper NLRs; they constitutively interact with each other regardless of activation^30–32^. Paired NLR genes can often be identified in the genome by their proximity to each other and divergent transcription^26^. Paired NLR proteins can be classified into ‘sensor’ NLRs and ‘executor/helper’ NLRs, with the sensor component often having an integrated domain (ID) that senses effector molecules^33^. The executor NLR often displays autoactivity upon overexpression, which can be suppressed by co-expression of the sensor NLR^30,34^. However, for some paired NLRs, expression of executor NLR without sensor NLR does not induce autoactivity^32^. The first NLR pairs reported to require each other for function were TIR-NLRs RPP2A and RPP2B^35^. Since then, many other paired NLRs, across dicots and monocot plant species, and of both TIR- and CC-NLR types, have been reported to function as pairs, such as RRS1/RPS4^36^, RRS1B/RPS4B^37^, CHS3/CSA1^38^, SOC3(WRR12)/CHS1^39^ TIR-NLRs, and RGA4/RGA5^31^ and Pik-1/Pik-2^40^ CC-NLRs^26^.

Arabidopsis RRS1-R (Resistance to *Ralstonia solanacearum* 1) and RPS4 (Resistance to *Pseudomonas syringae* 4) comprise a TIR-NLR pair^36,41^ that has been extensively studied as a model for paired NLRs (Fig. 1A). The RRS1-R allele from Ws-2 accession confers recognition of both AvrRps4 and PopP2, whereas the Col-0 RRS1-S allele lacks 83 amino acids at the C-terminus compared to RRS1-R and confers only AvrRps4 recognition. We previously reported the repression of RPS4 autoactivation by RRS1-R^30^, as well as de-repression of RRS1-R upon effector recognition^42^. RRS1-R and RPS4 recognize multiple effectors from diverse pathogen species; AvrRps4 from leaf pathogenic bacteria *Pseudomonas syringae pv. pisi*^43,44^, PopP2 from root pathogenic bacteria *Ralstonia solanacearum*^45–47^, XopJ6 from *Xanthomonas campestris* which causes black rot in *Brassicaceae*^48^, and unknown effector(s) from fungal pathogen *Colletotrichum higginsianum*^36^.

**Figure 1.**
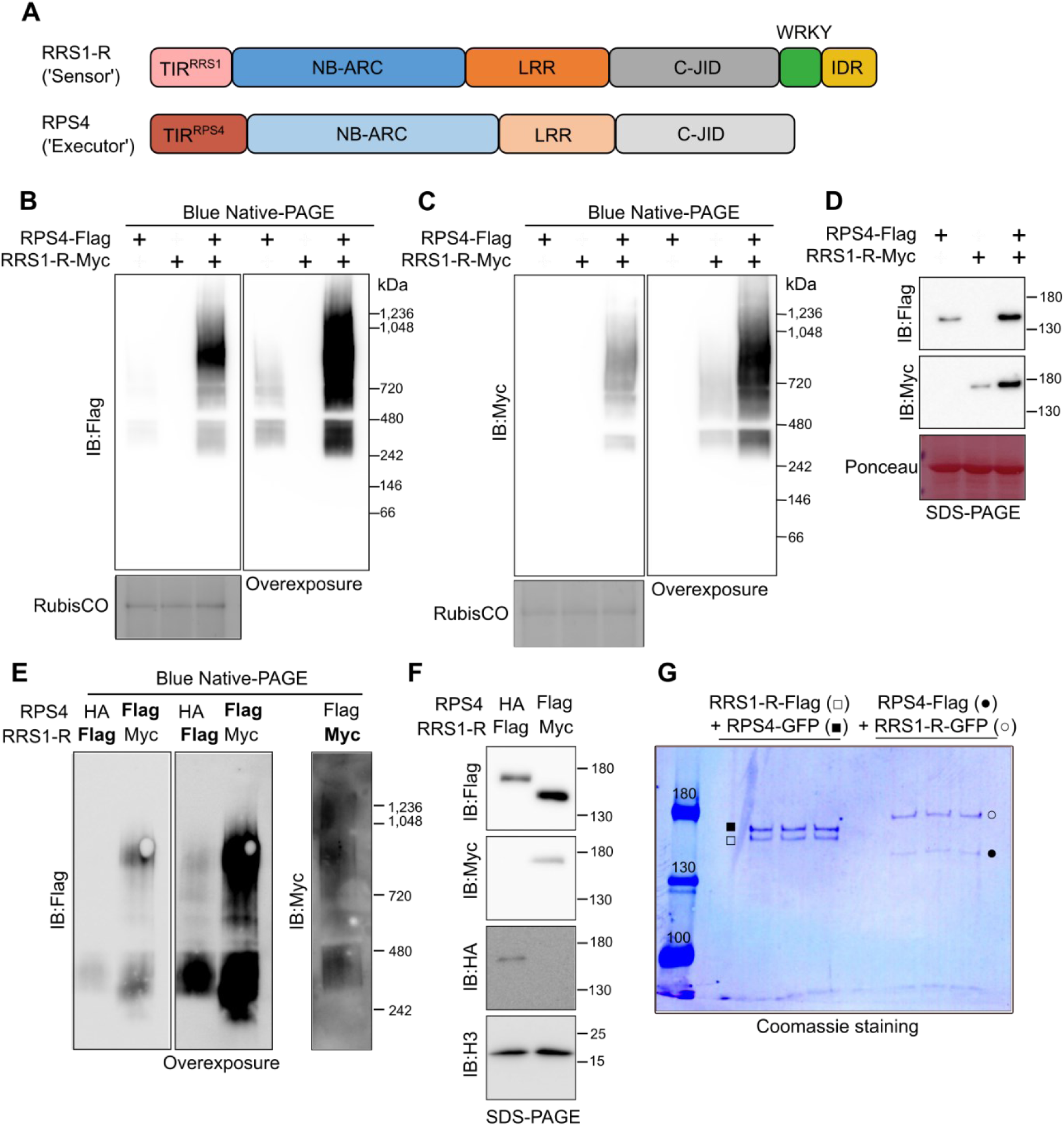
RRS1/RPS4 complex forms hetero-oligomeric complex in the inactive state. A. Schematic diagram of RRS1/RPS4 gene domains. B, C. RPS4 (B) and RRS1-R (C) in *Nicotiana benthamiana* (Nb) *eds1* show slower migration on blue native (BN)-PAGE. Transiently expressed RPS4-Flag and/or RRS1-R-Myc proteins were loaded on BN-PAGE, followed by immunoblotting. RubisCO complex serves as loading control. Molecular weights of markers are shown on the right. Overexposed images of the same membranes are indicated as ‘Overexposure’. Similar results were obtained at least three times. D. RRS1 and RPS4 proteins are stabilized upon co-expression. Samples from B and C were incubated in 3xSDS sample buffer with 100 mM DTT, loaded on SDS-PAGE, followed by immunoblotting. Molecular weights of markers are shown on the right. Ponceau S staining serves as loading control. Similar results were obtained at least three times. E. Slower migrating forms of RRS1-R and RPS4 are observed in Arabidopsis. Proteins extracted from Arabidopsis transgenic lines expressing RPS4-Flag with RRS1-R-Myc or RPS4-HA with RRS1-R-Flag (and inducible expression of AvrRps4) were loaded on BN-PAGE, followed by immunoblotting. Overexposed image of anti-Flag blotted membranes is indicated as ‘Overexposure’. Similar results were obtained at least three times. F. Protein accumulation of RRS1-R and RPS4 in Arabidopsis transgenic lines. Samples from E were incubated with SDS and DTT, loaded on SDS-PAGE, followed by immunoblotting. Immunoblots with anti-histone H3 antibody serve as loading control. Similar results were obtained at least three times. G. RRS1-R and RPS4 interact in 1:1 stoichiometry (Coomassie staining). Proteins from *N. benthamiana* plants transiently expressing RRS1-R-HF with YFP-RPS4 or RRS1-R-GFP with RPS4-HF. The protein extracts were immunoprecipitated using anti-Flag beads and eluted with 3xFlag peptide. The eluates were subsequently immunoprecipitated with anti-GFP beads, and bound protein complex were eluted with 3XSDS sample buffer. Each sample were loaded three times. Similar results were obtained in at least three biological replicates. Molecular weights of markers are shown on the left.

AvrRps4 is recognized by RRS1-R through interaction with the integrated WRKY domain^49^, while PopP2 and XopJ6 are acetyltransferases; the resulting WRKY acetylation activates the RRS1/RPS4 complex^50^. Modification of the WRKY domain of RRS1-R leads to an increase in intramolecular affinity between the RRS1-R TIR domain and the C-terminal IDR (Intrinsically Disordered Region) domain^51^. Effector recognition leads to changes in proximity between TIR domains of RRS1-R and RPS4 (TIR^RRS1^ and TIR^RPS4^, respectively) while maintaining RRS1/RPS4 interaction and protein abundance^51^.

Mimicking the cellular environment of proteins is crucial for studying their function. However, due to the diversity of proteins, and the sophisticated compartmentalization of cellular organelles, there is no “one size fits all” solution for plant protein purification^52^. In the current study, we needed to modify standard plant protein complex extraction protocols that typically include high levels of reducing agents (DTT) to alleviate browning of the extracts^34^. We found that these reducing agent conditions disrupted the complexes we were investigating and lower concentrations of DTT were essential for revealing protein complex properties because of a requirement for an oxidized cysteine at position 783 of RPS4.

We report here that the Arabidopsis RRS1-R/RPS4 complex does not change in size before or after effector recognition, and that the activated complex likely involves a reconfiguration that relieves TIR^RPS4^ domain inhibition by the TIR^RRS1^ domain. This enables a conformational change in the TIR^RPS4^ and TIR^RRS1^ domains that creates an NADase enzyme activity that we were able to measure. The TIR domain of ‘sensor’ NLR RRS1-R lacks the NADase active site as well as the aspartate residue in the BB loop, which is also lost in other sensor NLRs of the paired TIR-NLR type in Arabidopsis. Our study reports a novel mechanism of effector-dependent Arabidopsis paired TIR-NLR activation that ensures rapid immune activation upon effector recognition.

## Results

RRS1-R and RPS4 carry canonical TIR, NB-ARC and LRR domains, as well as additional domains (Fig. 1A). RPS4 carries a C-JID (C-terminal Jelly-roll/Ig-like domain), also known as PL (Post-LRR) domain, conserved in many TIR-NLRs^14,53^. Structural modelling using AlphaFold2 predicts that RRS1 also carries a C-JID, which in previous publications were reported as Domain 4^42^ (Fig. S1A-E). The C-JID of RRS1 partially aligns with that of RPP1, with additional α-helices predicted followed by the canonical C-JID (Fig. S1D-E).

### RRS1-R and RPS4 form a hetero-oligomer in *N. benthamiana* and *Arabidopsis*

To investigate native protein complex formation, we assessed protein migration in blue native-PAGE (BN-PAGE) after co-expression of RRS1-R and RPS4. RRS1-R-Myc (RRS1-R fused with 4xMyc tag) and RPS4-Flag (RPS4 fused with 3xFlag tag) were co-expressed transiently in *N. benthamiana eds1* mutant leaves and protein extracted from infiltrated leaves. Both proteins migrated in BN-PAGE at a size of ∼720 kDa (Fig. 1B, C). In the absence of its counterpart, both RPS4-Flag (Fig. 1B) and RRS1-R-Myc (Fig. 1C) migrated faster, with bands detected at ∼242 kDa and ∼480 kDa for both proteins. We concluded that both RRS1-R and RPS4 oligomerize with each other, consistent with reports from Williams *et al.* (2014)^30^. Huh *et al.* (2017)^34^ reported that RPS4 co-immunoprecipitates with itself only in the presence of RRS1^34^ using a different protein extraction condition from that used here. However, in our condition, RPS4 alone shows weak self-association (Fig. 1B), consistent with the auto-activity previously reported^34,54^. We also observed increased accumulation of both RRS1-R and RPS4 upon co-expression, suggesting they stabilize each other (Fig. 1D).

We performed BN-PAGE on protein extracted from two independent Arabidopsis transgenic lines (Fig. 1E) expressing RPS4-HA and RRS1-R-Flag with estradiol-inducible AvrRps4-expression^55^, and another Arabidopsis transgenic line overexpressing RPS4-Flag and RRS1-R-Myc (Fig. S1F, G). We detected both RRS1-R-Flag and RPS4-Flag from the two different transgenic lines in BN-PAGE. Both RRS1-R-Myc and RPS4-Flag migrated to above 720 kDa, and also below 480 kDa (Fig. 1E), consistent with our results using transient expression in *N. benthamiana*. RRS1-R and RPS4 thus form a complex in *N. benthamiana* indistinguishable from that formed in *Arabidopsis*. We also noted that RPS4-Flag forms below 480 kDa migrated faster than the corresponding RRS1-R-Flag forms (Fig. 1E), suggesting that these smaller forms may consist of RPS4 or RRS1-R only, respectively, considering that RPS4-Flag is smaller in size (∼140 kDa) compared to RRS1-R (∼160 kDa) (Fig. 1F). The extraction method enabled recovery of nuclear proteins, including histone H3, which is important since both RRS1-R and RPS4 reside in the nucleus^45,56^ (Fig. 1F).

### RRS1-R/RPS4 complex interact in 1:1 stoichiometry

We investigated the RRS1-R:RPS4 ratio in the complex by tandem-affinity purification of RRS1-R and RPS4 transiently co-expressed from *N. benthamiana*. Protein extracts were first immunoprecipitated with anti-Flag beads followed by elution with 3xFlag peptide and then immunoprecipitation with anti-GFP beads (Fig. S2A). The tandem-affinity purified samples were then visualized with Coomassie staining (Fig. 1G). Sequential affinity selection for each protein provides better information about the stoichiometry of the RRS1-R/RPS4 complex. RRS1-R and RPS4 interacted in approximately 1:1 ratio, which was also observed in samples reciprocally tagged with His-Flag (HF) tag and YFP/GFP tag to RRS1-R and RPS4 (Fig. 1G). Tandem affinity purification with different epitopes RPS4-Flag and RRS1-R-twinStrep transiently expressed in *N. benthamiana eds1* showed similar results (Fig. S2B). An RRS1-R/RPS4 heterotetramer is predicted to be ∼600kDa. We detected migration of an RRS1-R/RPS4 complex at ∼720 kDa (Fig. 1B, C), consistent with an RRS1-R/RPS4 complex that comprises two protomers each of RRS1-R and RPS4, with potential interactors additionally bound to the complex.

### The AE interface of TIR domain is required for RRS1-R/RPS4 hetero-oligomer formation

Previously, the crystal structure of TIR^RRS1^ and TIR^RPS4^ domains and their heterodimer revealed their structure consisting of five-stranded parallel β sheet (βA to βE) and five α-helical regions (αA to αE) positioned in tandem^30^. Importantly, this study revealed the AE interface (between αA and αE) is required for interaction (Fig. 2A)^30^, and TIR^RRS1^ and TIR^RPS4^ domains have higher affinity between their interfaces than either homodimer. This AE interface is also required to configure NADase activity^57^, while this AE interface-mediated interaction of TIR^RRS1^ and TIR^RPS4^ is crucial for repression of RPS4 TIR domains^30^. When we used the protein extracts from *N. benthamiana eds1* transiently expressing RRS1-R AE interface mutant (RRS1-R^AEmut^-MYC; alanine substitution of S25 and H26) with RPS4, we noticed significant reduction in RRS1-R/RPS4 complex on BN-PAGE (Fig. 2B). In the *eds1* background under moderate expression conditions, RPS4-Flag failed to co-immunoprecipitate RRS1-R^AEmut^-Myc (Fig. S3A). This indicates that the AE interface, originally identified as the binding interface of the TIR domains, plays an important role in the dimerization as well as the higher-order oligomerization of the full-length RRS1-R/RPS4 complex (Fig. 2B).

**Figure 2.**
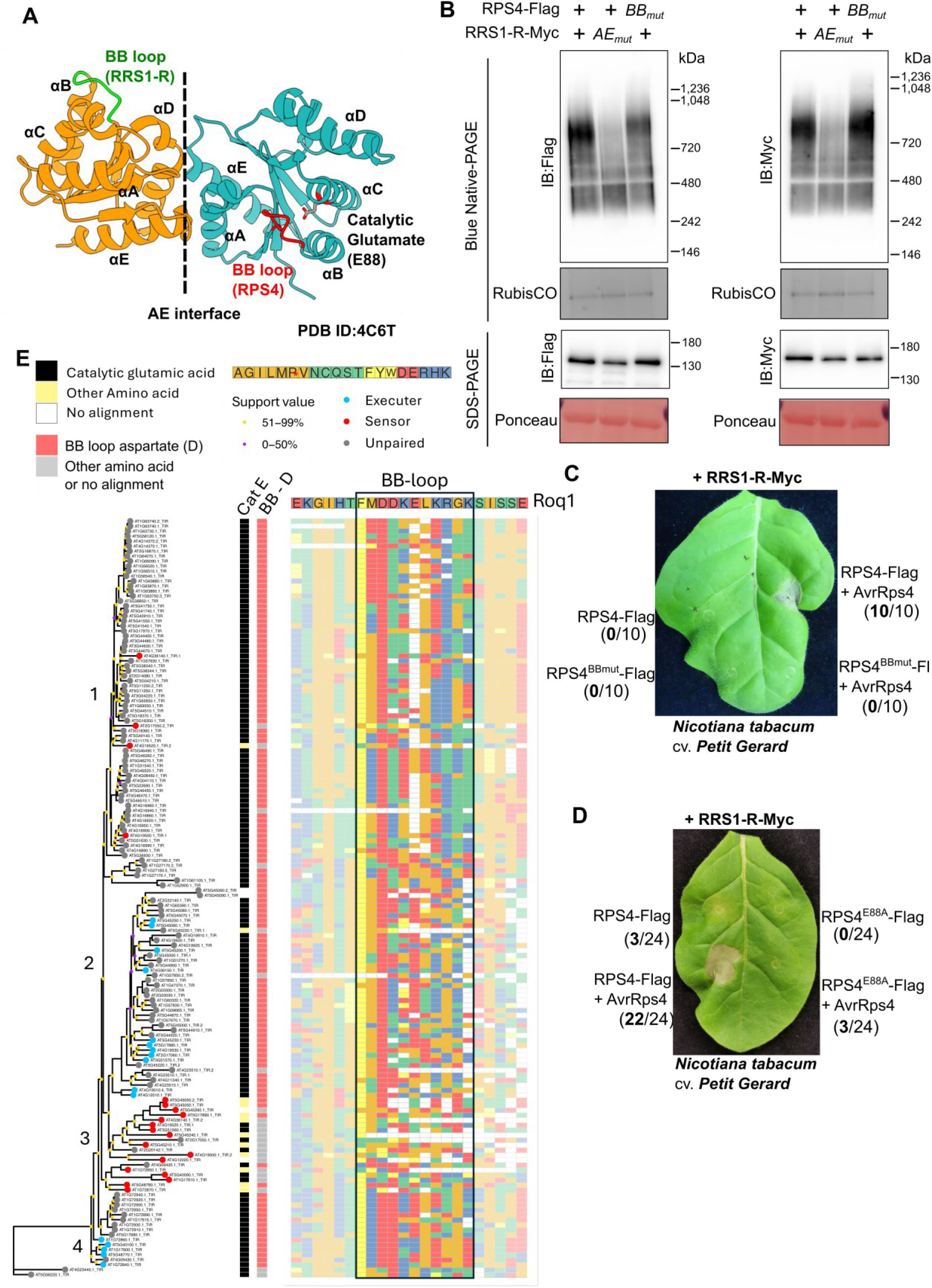
AE interface of RRS1-R TIR domain is required for RRS1/RPS4 hetero-oligomer formation, but NADase activity is not required for hetero-oligomerization. A. Structure of RRS1 and RPS4 TIR domain (PDB ID: 4C6T). AE interface (***dotted line***), and the relevant α helices are labelled. NADase active site (Catalytic glutamate, E88) of RPS4, BB-loop of RPS4 (red), and shorter BB-loop of RRS1 (green) are indicated. TIR^RRS1^ domains lack NADase active site. B. The AE interface of TIR^RRS1^ is essential for RRS1-R/RPS4 complex formation, while BB-loop is dispensable for RRS1-R/RPS4 complex formation. Proteins from *Nb eds1* leaves expressing wild-type or BB-loop mutation of RPS4-Flag (RPS4^BBmut^-Flag) and wild-type or AE interface mutant of RRS1-R-Myc (RRS1-R^AEmut^-Myc). The extracts were loaded on BN-PAGE or SDS-PAGE (after incubation with 3xSDS sample buffer and 100 mM DTT), followed by immunoblotting. RubisCO complex, and Ponceau S staining serves as loading control. Molecular weights of markers are shown on the right. C. Mutation of RPS4 BB-loop lacks effector-dependent activation by RRS1-R/RPS4. *Nicotiana tabacum* cv. *Petit Gerard* leaves expressing RRS1-R with RPS4 wild-type or RPS4 BB-loop mutation (RPS4^BBmut^-Flag) in the presence or absence of effector AvrRps4 were tested for Hypersensitive Response (HR). Images were taken 5 days post infiltration (dpi), and at least 3 leaves were used for each biological replicate. Number of HR occurrences (in bold) out of total number of leaves tested are indicated in parentheses. D. Mutation of RPS4 NADase active site (RPS4^E88A^) abrogates effector-dependent HR by RRS1-R/RPS4. *N. tabacum* cv. *Petit Gerard* leaves expressing RRS1-R with RPS4 wild-type or RPS4 NADase mutation (RPS4^E88A^-Flag) in the presence or absence of effector AvrRps4 were tested for HR. Images were taken 5 dpi, and at least 3 leaves were used for each replicate. Number of HR occurrences (in bold) out of total number of leaves tested are indicated in parentheses. E. Phylogenetic tree of TIR domains identified in Col-0 reference genome of Arabidopsis. 152 unique TIR domains from Col-0 reference genome were selected. Amino acids in the corresponding structural positions to the Roq1 catalytic glutamic acid (E86), conserved BB-loop aspartic acid (D45), the BB-loop (F43-K53) and surrounding region (E37-E58) were extracted from alignments and shown. The tree was rooted to divergent TIR-N proteins AT5G56220.1 and AT4G23440.1.

### BB-loop of RPS4 is required for HR upon effector recognition, but not oligomerization

Structural studies of TIR-NLRs Roq1 and RPP1 revealed that the BB loop (loop region connecting αB and βB) and the glutamate residue of the TIR domains (located in αC) are crucial for NADase activity (Fig. 2A)^18,19^. BB loop side of one TIR domain and the DE interface (between αD and αE) of another TIR domain create the NAD+ binding site^57^. The TIR^RPS4^ alone also requires its catalytic glutamate residue (E88) for HR^58^. We mutated the BB loop (RPS4^BBmut^; alanine substitution of I48, D49, and D50) or the NADase active site (RPS4^E88A^) of RPS4 and tested effector-dependent activation in *N. tabacum* cv. *Petit Gerard*. Both BB loop and NADase active sites of RPS4 are required for AvrRps4-dependent HR via the RRS1-R/RPS4 complex (Fig. 2C, D). However, the RPS4 BB-loop mutant (RPS4^BBmut^-Flag) co-expressed with RRS1-R still oligomerizes as does wild-type RPS4 (Fig. 2B). Similarly, the RPS4^E88A^ mutant is also unaltered in RRS1-R/RPS4 complex formation (Fig. S3B, C).

### BB-loop and NADase active site of sensor NLRs in a pair are degenerate

The TIR^RPS4^ has both the NADase active site and the BB loop, while TIR^RRS1-R^ and TIR^RRS1-S^ lack an NADase active site, and has a shorter BB loop, likely to abolish NADase activity^14^ (Fig. 2A). The TIR domains of the sensor NLR in other TIR-NLR pairs that are divergently transcribed have often lost the NADase active site, similar to RRS1-R and RRS1-S^19^. We identified 152 unique TIR domains from the Arabidopsis Col-0 proteome, and a phylogenetic tree was constructed. The structure of each TIR domain was predicted using AlphaFold2, and then aligned with TIR^Roq1^. The residue that aligns with catalytic glutamate (Cat E) of TIR^Roq1^, and the residue that aligns with aspartate residue (BB-D; third residue in the BB-loop) in the BB-loop of TIR^Roq1^, were visualized (Fig. 2E).

In most TIR domains, a conserved glutamate residue was identified at the catalytic site (Cat E), and conserved aspartate residue was identified at the BB-loop region (BB-D). However, in clade 3, there was lack of conservation in both the catalytic glutamate residue and aspartate residue in BB-loop (Fig. 2E). This clade includes many of the ‘sensor’ NLRs from paired TIR-NLRs found in Arabidopsis Col-0 (Table S3)^59^. Several TIR domains annotated to be sensor NLRs have two TIR domains. In all cases, one TIR domain was degenerate whereas the other TIR domain was functional^60^. All the functional domains were identified in clade 1. On the other hand, ‘executor’ NLRs of the paired TIR-NLRs were mostly distributed in the neighbouring clade 2 or clade 4 (Fig. 2E); they retain the catalytic glutamate and an aspartate in the BB-loop required for NADase activity.

This suggests that the sensor NLRs have evolved to repress the executor NLRs during the inactive state and this has selected for loss of NADase activity. It is currently unknown how this repression is relieved upon effector-dependent activation. However, the requirement of RRS1-R by autoactive RPS4^54^ suggest that, upon recognition of the effectors, these sensor NLRs then support the reconfiguration of the executor NLR for activation.

### P-loop of RPS4 but not RRS1-R is required for hetero-oligomer formation

The NB-ARC domain of plant NLRs is essential for oligomerization^5,27^. The P-loop motif binds phosphate groups of ATP or ADP, and is critical for NLR function^61^. We tested whether the P-loop motifs of RPS4 and RRS1-R NB-ARC domains are required for oligomerization. RPS4 P-loop mutant (RPS4^K242A^) lost effector-dependent activation, whereas RRS1-R P-loop mutant (RRS1-R^K185A^) did not^30^, irrespective of epitope tags fused to RRS1-R and RPS4 (Fig. S4A).

We observed that the RPS4^K242A^ mutant was unable to oligomerize into an RRS1-R/RPS4 complex, similar to the AE interface mutant (Fig. S3B, S4B). The RRS1-R^K185A^ mutant was able to form RRS1-R/RPS4 oligomer, but not as much as the wild-type (Fig. S4B). This suggests that RRS1-R may not require nucleotide binding, whereas nucleotide binding is crucial for RPS4 to form the RRS1-R/RPS4 complex, and this correlates with failure of RPS4^K242A^ to activate immune response upon AvrRps4 recognition.

### RPS4 C783 is required for RRS1-R/RPS4 oligomerization

We noticed that RRS1-R/RPS4 complex dissociated at high concentrations of DTT (10 mM; Fig. S5A). This prompted us to test whether disulfide bonds were formed within the RRS1-R/RPS4 complex, either as intramolecular bonds, or as intermolecular bonds between RRS1-R and RPS4. We noticed that slower-migrating form of RPS4 and RRS1-R may be due to oxidation occurring post-extraction (Fig. S5B, C). To prevent oxidation of protein extracts generating artefactual disulfide bonds during extraction, we added N-ethylmaleimide (NEM) to the protein extract to prevent free sulfhydryl groups from forming disulfide bonds *in vitro* (Fig. S5D). After purification of proteins from *N. benthamiana*, multiple disulfide bonds were identified through LC-MS/MS (Fig. S5E, F). The three main disulfide linkages identified from RPS4 and RRS1-R were intramolecular disulfide bonds, RPS4^C783^-RPS4^C1020^, RPS4^C1001^-RPS4^C1006^, and RRS1-R^C289^-RRS1-R^C291^.

However, only one of the identified cysteine residues, C783 of RPS4, was required for HR upon AvrRps4 recognition (Fig. S6A, B). Unexpectedly, RPS4^C1020^ did not affect effector-dependent activation (Fig. S6B). RPS4^C783A^ was lower in abundance compared to wild-type RPS4 (Fig. S6E). However, when the samples were loaded on blue native-PAGE (Fig. S6C, D), slow-migrating form corresponding to RRS1-R/RPS4 complex was not observed with RPS4^C783A^, and smearing patterns were observed instead. This suggests that RPS4^C783A^ can co-immunoprecipitate RRS1-R (Fig. S6E), but cannot assemble into wild-type complexes and form aggregates (Fig. S6C, D).

The C783Y mutation of RPS4 was previously identified in mutant screens for suppressors of the RRS1-R^slh1^ autoactivity (Fig. 3A)^62^. C783 resides within the LRR motif with consensus sequence (LxxLxxLxxLxLxx(**N/C/T**)x(x)LxxIPx)^63^. We introduced serine substitution of C783 (RPS4^C783S^), to maintain the polarity of the cysteine residue. Replacement of C783 to serine also resulted in loss of effector recognition (Fig. 3B). When protein extracts expressing RPS4^C783S^-Flag and RRS1-R-Myc were loaded on BN-PAGE, we observed loss of oligomeric RRS1-R/RPS4 complex (Fig. 3C), both in the presence and absence of AvrRps4. We also observed reduction of RPS4 protein accumulation (Fig. 3D), suggesting RPS4 protein stability may also be affected by C783 mutation. Thus, C783 of RPS4 is required for RRS1-R/RPS4 complex oligomerization as well as stabilization of the RPS4 protein.

**Figure 3.**
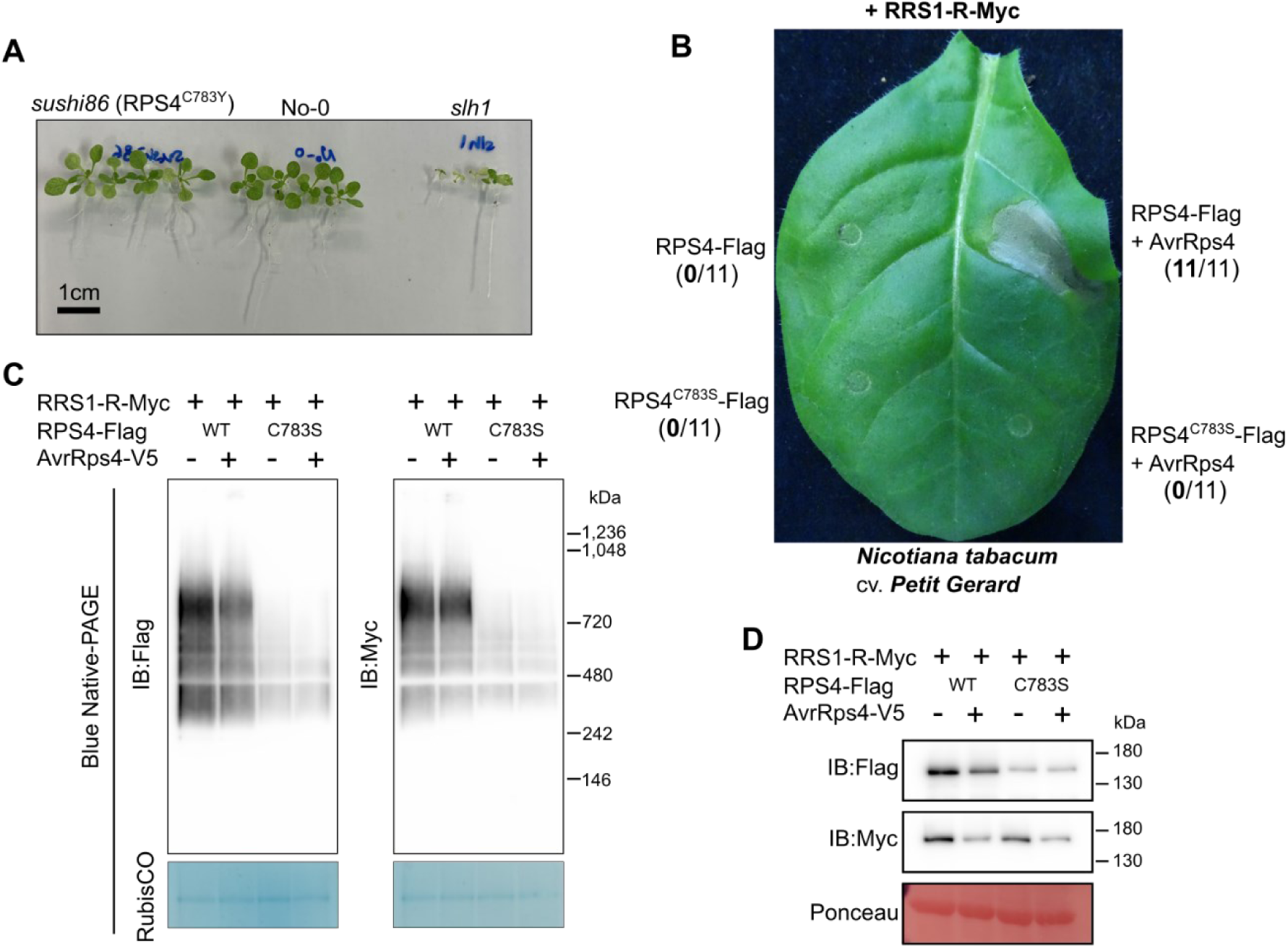
C783 of RPS4 is required for RRS1-R mediated autoactivity as well as effector-dependent activation. A. RPS4 C783Y (*sushi86*) suppress autoactive *slh1* phenotype. Arabidopsis *sushi86*, No-0 and *slh1* seedlings were germinated on ½ MS media with 1% sucrose and grown for 10 days. B. C783S mutation of RPS4 lost RRS1-R-dependent, effector-dependent HR. *N. tabacum* cv. *Petit Gerard* expressingRPS4 wild-type or mutant with RRS1-R in the presence or absence of effector AvrRps4 in *N. tabacum* cv. *Petit Gerard* were tested for HR. Images were taken at 5 dpi, and at least 3 biological replicates were performed. Number of HR occurrences (in bold) out of total number of leaves tested are indicated in parentheses. C. Slower migrating forms of RRS1-R and RPS4 are lost in RPS4^C783S^ mutant. Proteins were extracted from *N. benthamiana eds1* leaves expressing RPS4-Flag and its RPS4^C783S^ mutant with RRS1-R-Myc in the presence or absence of effector AvrRps4. Extracts were loaded on BN-PAGE, followed by immunoblotting. RubisCO complex serves as loading control. Molecular weights of markers are shown on the right. D. Protein accumulation of RPS4^C783S^ is weaker than wild-type RPS4. Extracts from C were incubated in 3xSDS sample buffer with 100 mM DTT and loaded on SDS-PAGE, followed by immunoblotting. Ponceau S staining of the SDS-PAGE serve as loading control. Molecular weights of markers are shown on the right.

### Activation of RRS1-R/RPS4 does not alter migration in non-denaturing gels in *N. benthamiana* and Arabidopsis

To assess changes in oligomerization of the RRS1-R/RPS4 complex upon effector-dependent activation, singleton TIR-NLR Roq1, and RRS1-R/RPS4 were co-expressed with their cognate effectors XopQ or PopP2, respectively, in *N. benthamiana eds1*. While with Roq1 we observed induced oligomerization upon activation, we did not observe changes in the size of RRS1-R/RPS4 complex in the presence of PopP2 (Fig. 4A). This suggests that small conformational changes by the effector are sufficient to trigger activation of RRS1-R/RPS4 and that activation does not require further oligomerization. Although we detected reduction of Roq1 protein accumulation upon activation, there were no significant changes in RRS1-R or RPS4 protein accumulation (Fig. 4B).

**Figure 4.**
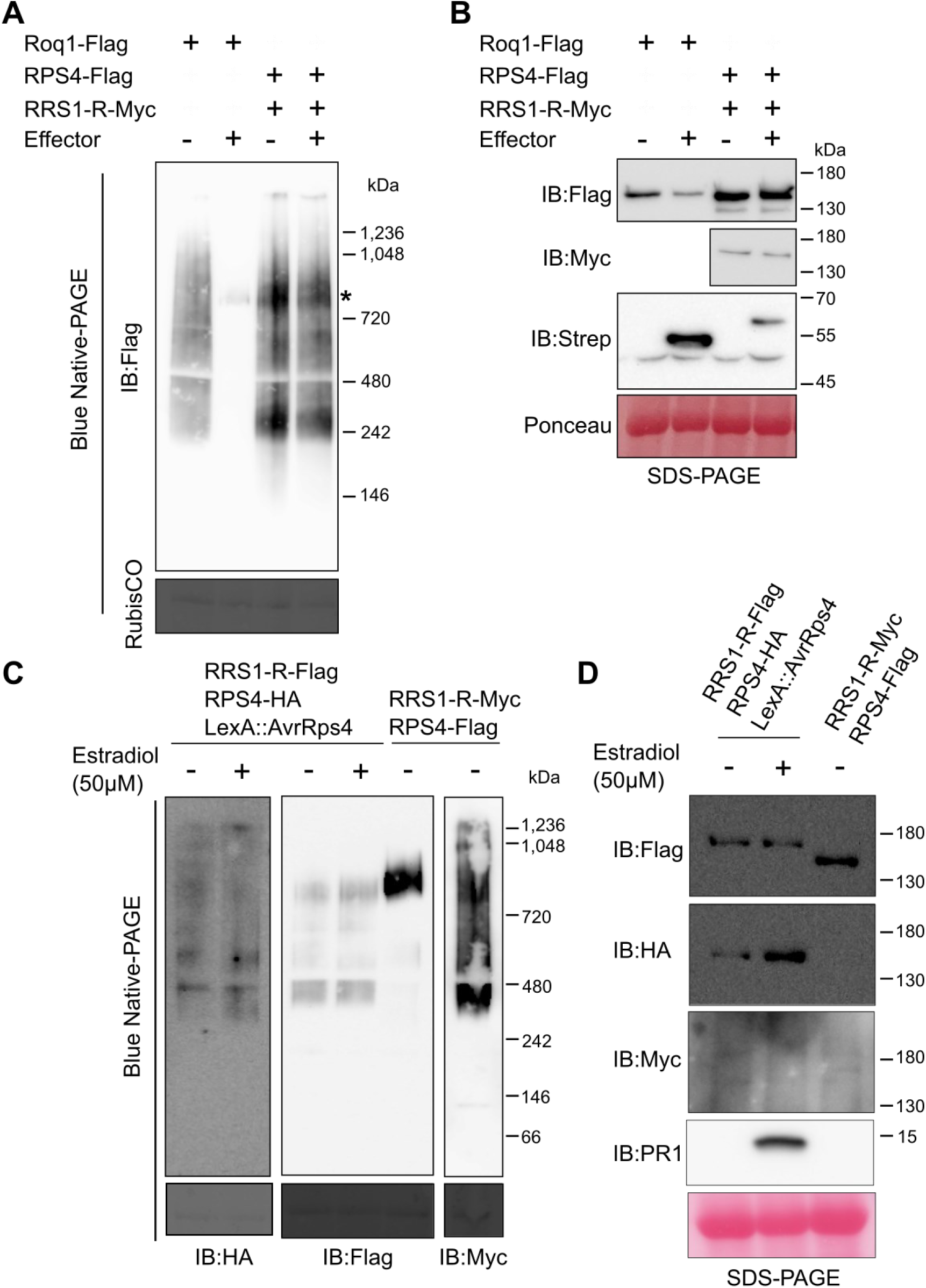
Activation of RRS1/RPS4 does not alter complex size. A. RRS1-R/RPS4 complex does not alter complex size upon activation by cognate effector (Strep-XopQ for Roq1, and PopP2-Strep for RRS1-R/RPS4), in contrast to activated singleton TIR-NLR Roq1. Proteins were extracted from *N. benthamiana eds1* expressing inactive and active forms of Roq1 and RRS1/RPS4, loaded on BN-PAGE, followed by immunoblotting. Molecular weights of markers are shown on the right. Similar results were obtained at least three times. B. Samples from A were incubated with 3xSDS sample buffer and 100 mM DTT at 70°C, and loaded on SDS-PAGE, followed by immunoblotting. Ponceau S staining serve as loading control. Molecular weights of markers are shown on the right. Similar results were obtained at least three times. C. Induced expression of AvrRps4 in Arabidopsis transgenic line does not alter RRS1/RPS4 hetero-oligomer size. Arabidopsis transgenic lines expressing RPS4-Flag with RRS1-R-Myc or RPS4-HA with RRS1-R-Flag with estradiol-inducible AvrRps4, loaded on BN-PAGE, followed by immunoblotting. Molecular weights of markers are shown on the right. Similar results were obtained at least three times. D. PR1 is induced upon estradiol-inducible AvrRps4 expression. Samples from C were incubated with 3xSDS sample buffer and 100 mM DTT, loaded on SDS-PAGE, followed by immunoblotting. PR1 expression was checked with anti-PR1 antibody. Ponceau S staining serve as loading control. Molecular weights of markers are shown on the right.

We also used the AvrRps4-inducible Arabidopsis transgenic line^55^ to compare inactive and active forms of the RRS1-R/RPS4 complex. We observed no difference in migration patterns in the presence or absence of effector AvrRps4 induction (Fig. 4C). We detected PR1 protein accumulation as control, induced upon expression of AvrRps4^55^, which was only observed in the AvrRps4-induced samples (Fig. 4D). This further suggested that the RRS1-R/RPS4 complex does not undergo changes in complex size upon activation and likely remains as a heterotetramer.

### NADase activity is only observed in activated RRS1-R/RPS4, and enzyme activity is not required for hetero-oligomer formation

Next, we tested for hydrolysis of εNAD^+^ with purified RRS1-R/RPS4 complex expressed in *N. benthamiana eds1* plants. RRS1-R/RPS4 complex was purified using Anti-Flag agarose beads, and eluted with 3xFlag peptide (Fig. 5A). εNAD^+^ is an analog of NAD+ that upon cleavage by NAD+ hydrolases, produces fluorescent εADP-ribose product^64^. We observed an increase in fluorescence resulting from hydrolysis of εNAD^+^ only from the active RRS1-R/RPS4 complexes that were co-expressed with cognate effector PopP2, and minimal changes in fluorescence from the inactive complex or without εNAD+ substrates (Fig. 5B). Adding ATP did not make any difference in the NAD+ cleavage, suggesting that the activated RRS1-R/RPS4 complex does not use ATP as substrate. The purified RRS1-R/RPS4 complex loaded on BN-PAGE mainly migrated as ∼242 kDa protein complexes, and we observed more RRS1-R/RPS4 complexes around ∼720 kDa in its activated state (Fig. 5C). Conceivably, this is due to the increased stability of the RRS1-R/RPS4 complex upon effector recognition.

**Figure 5.**
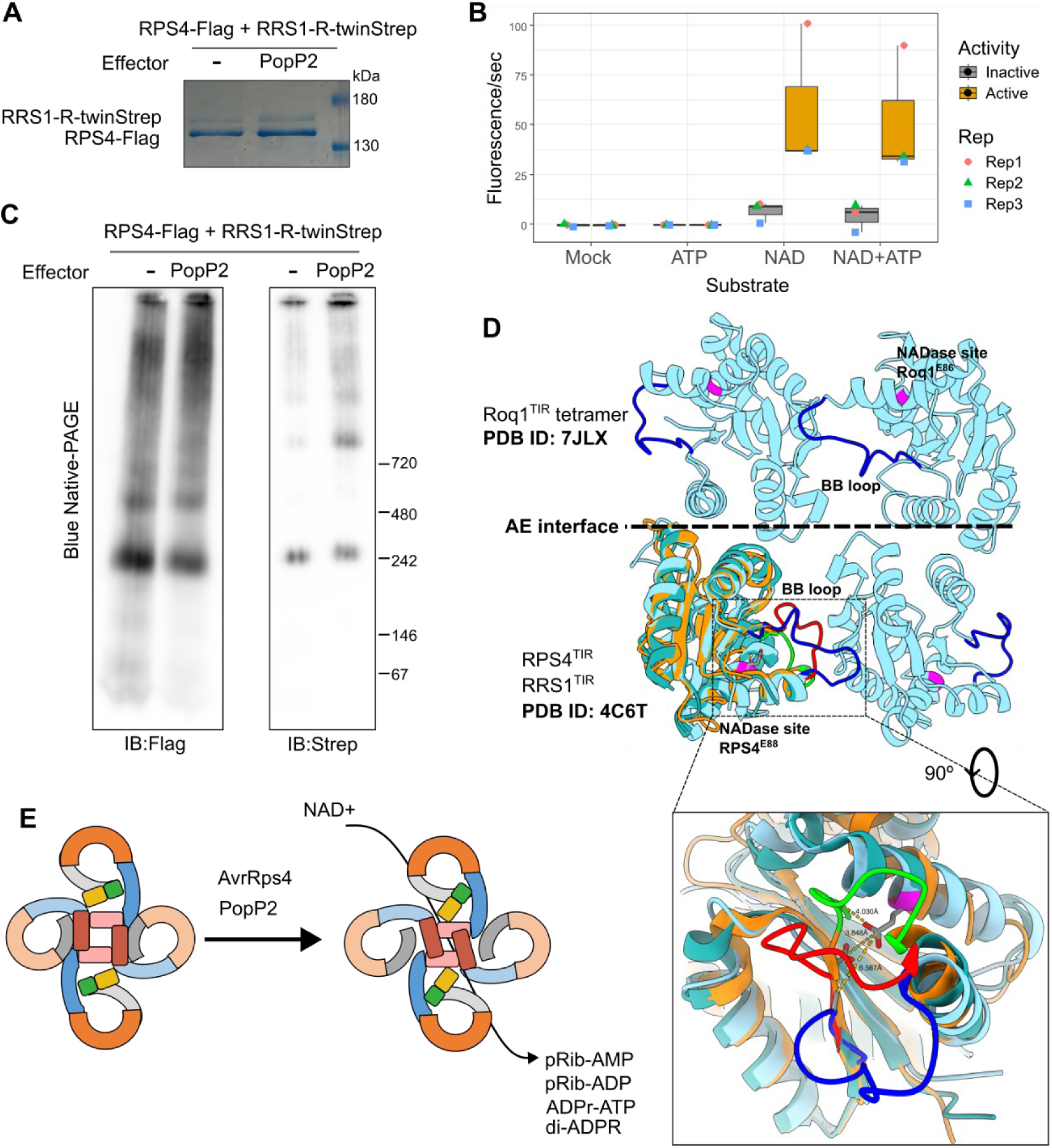
A. Purified RRS1-R/RPS4 complex. RPS4-Flag and RRS1-R-twinStrep were co-expressed with or without effector PopP2 in *N. benthamiana eds1* and purified using anti-Flag beads. Immunoprecipitated proteins were eluted using 3xFlag peptide, denatured with SDS and DTT, then subsequently loaded on SDS-PAGE for Coomassie staining. Molecular weights of markers are shown on the right. Similar results were obtained at least three times. B. NADase activity of purified, active RRS1/RPS4 complex. Purified RRS1-R/RPS4 complex from A were incubated with substrates. Fluorescence emission by ε-NAD cleavage is measured in time-course of 1 min for 1 h, and reaction rate was plotted as Fluorescence/sec. C. Purified RRS1-R-/RPS4 protein complex from A were loaded on BN-PAGE, followed by immunoblotting. Molecular weights of markers are shown on the right. Similar results were obtained at least three times. D. Small conformational change of the BB-loop may be sufficient to drive NADase activity activation of the RRS1-R/RPS4 complex. TIR domain of RRS1-R and RPS4 (RRS1-R, orange; RPS4, blue), and TIR domain of activated Roq1 with open BB-loop (light blue) is mapped together. Inset shows magnified view of the BB-loop region and the distance of the core C of aspartate (D) in the BB-loop to the catalytic glutamate (E88 for RPS4). E. Schematic diagram for activation mechanism of RRS1/RPS4 complex. RRS1-R/RPS4 complex is present as heterotetramer in the inactive state. Upon modification of the WRKY domain by AvrRps4 or PopP2, the RRS1-R/RPS4 complex undergo conformational change that leads to activation of NADase activity.

### Small conformational changes likely activate the RRS1/RPS4 complex

We compared the crystal structure of TIR^RRS1^ and TIR^RPS4^ domains^30^ with the cryo-EM structure of TIR^Roq1^ in the activated Roq1 resisotosome^15^. In the ‘dimer of dimer’ configuration of Roq1 TIR domains, the BB-loop of one TIR domain moves away from the NADase active site, while the other TIR domain provides the DE interface. This results in the two TIR domains creating the BE interface where NAD+ binds to (Fig. 5D). Compared to this, both TIR^RRS1^ and TIR^RPS4^ BB-loops are positioned closer to the NADase site of TIR^RPS4^ (RPS4^E88^), while the overall structure of RRS1-R and RPS4 TIR domains are in alignment with Roq1. This strongly suggests that only a minor change in configuration of the heterotetrameric RRS1-R/RPS4 complex is required for activation to occur.

To test the feasibility of the RRS1-R/RPS4 heterotetramer, we deployed AlphaFold to predict models of RRS1-R/RPS4 TIR domains (Fig. S7). Compared to the tetrameric TIR domain structure of Roq1 (Fig. S6A), TIR^RRS1^ and TIR^RPS4^ alone in the absence of NAD+ do not appear to model into a ROQ1-like tetramer (Fig. S7D, F). When modelling two TIR^RPS4^ and two TIR^RRS1^ together, 4 out of 5 predicted models assume the ROQ1-like conformations (Fig. S7B), which suggests that tetrameric configuration of two molecules of TIR^RPS4^ and two molecules of TIR^RRS1^ can be accommodated in the inactive state. This arrangement has the hall-mark AE interface intact, and also the BE interface^57^, with RPS4 contributing the BB-loop side, and RRS1-R contributing the DE surface side of this interface.

Addition of NAD+ in the predictions did not improve the confidence of the TIR^RRS1^ models (Fig. S7G) but dramatically improved the confidence in the RPS4 tetramer models (Fig. S7E). This also supports the possibility that RPS4 may form a homotetrameric resistosome. All five models assume a ROQ1-like conformation, with high confidence. Addition of NAD+ did not change the confidence in the overall RRS1-R/RPS4 models, but disrupts the formation of BE interfaces in the prediction. How this conformational change is induced, especially by rearrangement of the other domains of RRS1-R and RPS4 proteins will need to be resolved through structural analyses.

In summary, we report that Arabidopsis paired NLRs RRS1-R/RPS4 complex is in a ‘poised’ state as a heterotetramer, which upon recognition of its cognate effector, maintains its stoichiometry but alters its conformation to induce NADase activity at the TIR domains (Fig. 5E).

## Discussion

Paired NLRs are found in diverse plant species, and play important roles in disease resistance in crops^26^. Transfer of such paired NLRs together between species can elevate resistance to disease ^65^. Such paired NLRs can also provide a chassis to create novel disease resistance^66–71^. Despite the remarkable progress in structural analysis of activated NLRs, we still have much to learn about paired NLR immune receptors. The diversity of paired NLRs, and the distinction between paired TIR-NLRs and paired CC-NLRs, suggest the potential for mechanistic differences between paired NLRs.

### Multiple domain-domain interactions are essential for constitutive protein complex assembly of RRS1-R/RPS4

Which interfaces of each NLR are involved in complex formation? We observed that mutations in each of the TIR (Fig. 2B), NB-ARC (Fig. S3B), and LRR (Fig. 3C) domains of RRS1-R and RPS4 can compromise formation of an RRS1-R/RPS4 complex. This is in contrast to a previous report using different constructs^30^ suggesting that full-length RRS1-R and RPS4 interact even with TIR domain AE interface mutations. However, we now find using new constructs with different tags, and with lower expression, and different protein extraction conditions, that RRS1-R-Myc AE interface mutants are not co-immunoprecipitated by RPS4-Flag wild-type proteins (Fig. S3A). This is consistent with the results obtained from another paired TIR-NLR CHS3/CSA1^72^. and suggests TIR^RPS4^-TIR^RRS1^ interactions are essential for complex formation.

The P-loop of RPS4, but not RRS1-R, is required for defense activation^30^. We show here that the P-loop of RPS4 is required for oligomer formation, but the P-loop of RRS1-R is not (Fig. S4). The P-loop, or Walker A motif is required for binding to the β- and γ-phosphates of the bound NTP^5^. Thus, mutation of the P-loop motif would result in lack of ADP/ATP binding in the pocket. Interestingly, based on analysis from Ma *et al.* (2020)^14^, RPS4 is likely to preferentially bind ADP due to the S346 residue interfering with the position of the γ-phosphate group, whereas RRS1-R may preferentially bind ATP^27^. However, our results suggest that RRS1-R may not bind to nucleotides, or binding of nucleotides is unnecessary (Fig. S4). RRS1-S can suppress the action of RRS1-R, explaining why RRS1-R is recessive^54^; we previously reported that this suppression requires the P-loop of RRS1-R^30^, and this remains unexplained.

The LRR domain and C-JID domain of TIR-NLRs function in recognizing effectors^14,15^. We have previously identified multiple residues along the LRR and C-JID domains of RPS4, including C783, that are required for autoactivity of the RRS1-R^slh1^ mutant^62^. We show here that the C783 residue of RPS4 in the LRR domain is required for stability and oligomerization of the RRS1-R/RPS4 complex (Fig. 3C). C783 is positioned within the canonical motif of the LRR repeat^63^, suggesting C783 may have structural roles in maintaining the LRR domain in a configuration that enables constitutive interaction with RRS1-R.

### The oligomer of RRS1-R/RPS4 complex is detected under low DTT concentrations

The outcomes of biochemical analyses of proteins are highly dependent upon extraction and buffer condition^52^. We detected a stable RRS1-R/RPS4 complex when purified under low-reducing conditions (Fig. S5A) that is not seen in standard 10 mM DTT conditions used previously^34^. This implies cysteine residues in RRS1-R or RPS4 may be involved in forming disulfide bonds. Indeed, in our mass spectrometry dataset, we identified intramolecular disulfide bonds of RPS4, which also included C783-C1020 of RPS4 (Fig. S5C). However, mutating the C1020 residue of RPS4 did not disrupt the effector recognition (Fig. S5H). Recent reports have shown sulfenylation of proteins due to oxidation^73^. Sulfenylation can render the cysteine residues prone to forming disulfide bonds with neighbouring cysteine residues^74^. Thus, alternative modifications other than disulfide bonds may occur in C783 of RPS4, that is subsequently modified to form disulfide bonds. Consistent with this prediction, C783 forms disulfide bonds with multiple cysteine residues of RPS4 (Table S4).

### Why has the sensor NLR in a paired TIR-NLR lost NADase activity?

We show here that there are no changes in size of the RRS1-R/RPS4 complex in the presence or absence of cognate effector. This is consistent with previous studies on RRS1-R and RPS4 interactions that show no apparent change upon co-expression with a cognate effector^34,55^ while displaying changes in proximity of TIR^RPS4^ and TIR^RRS1^ upon effector recognition^51^. This constitutive interaction between RRS1-R and RPS4 may have selected for loss of NADase activity in the RRS1-R sensor, reducing the risk of constitutive NADase activity. The sensor NLRs of most paired TIR-NLRs have degenerate BB loops and lack the NADase active site (Fig. 2E), in addition to their genetic linkage to an executor TIR-NLR and the presence of ID. This feature may help identify novel paired TIR-NLRs in different plant genomes, especially those that have sensor NLRs without an ID^75^.

This evolution of paired TIR-NLRs can be explained as a unidirectional evolutionary ‘ratchet’^76^. The proximal positioning of the paired TIR-NLRs in the genome, as well as the similarity of their NB domains^59^ imply a possible duplication event. The constitutive interaction between the paired TIR-NLR proteins would have reinforced shortening of BB loop, to prevent constitutive NADase activity, while NADase activity from executor NLR alone did not affect the function of the overall paired TIR-NLR complex. However, due to these mutations, the sensor NLRs could no longer function on their own, and strictly require the executor NLRs for function.

### How does the RPS4/RRS1 immune receptor work?

RRS1-R represses autoactivation of RPS4 but is also required for effector-dependent activation of the RRS1-R/RPS4 paired NLR. Roq1 and RPP1 resistosomes assemble into a ‘dimer of dimer’ configuration of TIR domain that generates 2 NADase active sites in one resistosome^14,15^. Only one of the TIR domains undergoes conformational change at the BB loop that reveals the NADase active site, while the other TIR domain supports the NADase activity via creating the BE interface. Conceivably, the NADase active site and BB-loop of RPS4 in the “dimer of a heterodimer” of RRS1-R/RPS4 complex are sufficient for activation. Thus, the function of sensor NLRs goes beyond their role of repressing executor NLRs and also includes providing the substrate binding interface upon activation.

Given the auto-activity of RPS4 alone, RPS4 might also be able to form a homotetramer, which may be stabilized in the presence of NAD+ as shown in the model prediction (Fig. S6E). Nonetheless, RPS4 itself shows weak autoactivity compared to effector-dependent, RRS1-R-dependent activation, due to weak affinity in self-association (Fig. 1B)^30,34,53^. We previously reported that the HR conferred by autoactive alleles of RPS4 require co-expression with RRS1-R; RRS1-S does not suffice^54^. This strongly suggests additional role of RRS1-R during activation, possibly by providing the BE interface for NAD+ binding. Thus, the constitutive interaction of RPS4 and RRS1-R evolved into TIR^RPS4^ domain providing the NADase active site, and TIR^RRS1^ domain supporting RPS4 by effector-dependent reconfiguration of the complex to relieve inhibition of TIR^RPS4^ by TIR^RRS1^, enabling active TIR^RPS4^ conformation to form and create the NAD+ binding site. The strong affinity between TIR^RPS4^ and TIR^RRS1^ domains are consistent with this hypothesis, as well as the weak autoactivity of RPS4 and strong autoactivity of RPS4 autoactive allele with RRS1-R^30,53,54^.

Previous structural studies have reported direct binding of the WRKY domain of RRS1-R with PopP2 and AvrRps4^49,50^. However, we could not detect changes in the migration of RRS1-R/RPS4 complex in the BN-PAGE analysis in the presence or absence of these effectors. This is consistent with previous studies showing a low affinity of PopP2 for RRS1-R/RPS4^30,34^. In addition, the small size of truncated AvrRps4^44^ may not change apparent size of active RRS1-R/RPS4 protein complex as assayed in BN-PAGE gels. Domain-domain interaction studies of RRS1-R and RPS4 have shown that the C-terminal domains undergo a large conformational change upon effector recognition^42,51^. Conceivably, in the full-length context of the RRS1-R/RPS4 complex, large conformational changes eject the bound effectors from the activated RRS1/RPS4 complex. Further structural studies may reveal these intermediary states of RRS1-R/RPS4 complex before and after activation.

In conclusion, our analyses suggest that the Arabidopsis paired NLR RRS1-R/RPS4 complex is in a ‘poised’ state as a dimer of a heterodimer prior to effector recognition. Another report regarding the Arabidopsis paired NLR CHS3/CSA1 showed oligomerization into a heterotetramer^72^ but was not able to address recognition-dependent changes. We present here a mechanism whereby NLRs can rapidly respond to ligand perception without altering stoichiometry or forming new protein complexes.

## Acknowledgements

Authors wish to thank the past and present members of the Jones lab, and particularly Wen Huang for technical support. We thank Tiancong Qi (Tsinghua University), Brian Staskawicz (University of California Berkeley), Yejin Ahn and Kee Hoon Sohn (Seoul National University) for sharing their materials. H.-K.A. wishes to thank Timothy Wells, John Cotton (John Innes Centre horticultural services) for the care of plants; Mark Youles, Liam Egan for providing the Golden Gate modules for cloning; Aleksandra Wawryk-Khamdavong, Jodie Taylor, and Matthew Smoker for transformation of Arabidopsis plants used in this study. We would also like to thank group of Jijie Chai at West Lake University for helpful discussion.

## Funding information

ERC Advanced grant “ImmunityByPairDesign (IBPD; ref. 669926)” to H.-K.A., N.M., M.J.B and J.D.G.J., UKRI-BBSRC for Plant Health ISP (BB/P012574 and BBS/E/J/000PR9795), and Advancing Plant Health ISP (BB/X010996/1) to M.J.B and J.D.G.J., UKRI-BBSRC grant (BBS/E/J/000PR9797) to M.T.H. and W.M., UKRI-BBSRC NRPDTP to L.K., Royal Society University Research Fellowship (URF\R1\241258) to H.-K.A., and Gatsby Charitable Foundation funding to H.-K.A., S.P.Y.K., S.U.H., J.S., M.T.H., M.S., J.C., F.L.H.M, W.M., and J.D.G.J. G.G. was supported by the special project of long-term overseas training for young scientific and technological talents in 2022 of CAS and the Major Project of Agricultural Biological Breeding (2023ZD0402502).

## Author Contributions

Conceptualization, H.-.K.A., J.D.G.J.; Methodology, H.-.K.A., J.S., N.M.; Validation, H.-K.A., G.G., S.P.Y.K.; Formal analysis: H.-.K.A., M.T.H.; Investigation: H.-.K.A., G.G., S.P.Y.K., S.U.H., J.S., M.T.H., H.B., L.K., H.Z., N.M., M.S., J.C.; Resources, J.D.G.J.; Data Curation: J.S., Writing – Original Draft: H.-.K.A., J.D.G.J.; Writing – Review & Editing: all authors, Visualization: H-K.A., G.G., S.P.Y.K., M.T.H., H.B.; Supervision: H.-K.A., M.J.B., F.L.H.M., W.M., J.D.G.J.; Funding Acquisition: H.-.K.A., F.L.H.M., M.J.B., W.M., J.D.G.J.

## Declaration of Interests

The authors declare no competing interests.

## Materials and Methods

### Plant materials

*Nicotiana benthamiana* wild-type and *eds1* plants ^77^, and *Nicotiana tabacum* cv. Petit Gerard plants were grown in long-day conditions (16 h light/8 h dark) at 22°C, humidity 45-65%. Arabidopsis transgenic lines (Table S2) were grown in short-day conditions (8 h light/16 h dark) at 22°C for 4-5 weeks, then moved to long-day conditions for seed propagation. Transgenic Arabidopsis line seedlings were grown in ½ MS liquid culture with 1% sucrose for 7 days under constant light at 22°C.

### Golden Gate cloning

To generate constructs of epitope tag-fused RRS1-R or RPS4 proteins, Golden Gate cloning ^78^ was used. Generated constructs are listed in the Supplemental table 1. All the modules were provided by TSL Synbio team.

### Agrobacterium infiltration

Binary constructs for transient expression were transformed into *Agrobacterium tumefaciens* strain GV3101-pMP90. Agrobacterium cells were resuspended in infiltration buffer (10 mM MgCl_2_ and 10 mM MES, pH 5.6, 100 μM acetosyringone). OD600 of each construct-containing Agrobacterium was adjusted to 0.5, and OD600 for Agrobacterium containing p19 was adjusted to 0.2. Infiltration was performed after 1-2 h incubation at room temperature. The infiltrated *N. benthamiana* leaves were harvested 3 days post infiltration (3 dpi).

### Hypersensitive Response (HR) assay

As described, Agrobacterium cells harboring the constructs were resuspended in infiltration buffer and subsequently infiltrated into 4 to 5 week-old *N. tabacum* plants. The leaves were imaged at 5 dpi.

### Protein extraction for blue-native PAGE analysis

Protein extraction was performed as described in Ahn, Lin *et al.* (2023) ^27^ with minor adjustments. Agrobacterium-infiltrated *N. benthamiana* leaves or Arabidopsis seedlings/rosette leaves were harvested, and immediately frozen in liquid nitrogen. Frozen samples were then ground with Geno/Grinder® (SPEX SamplePrep). Protein extraction buffer (50 mM Tris-Cl pH 7.5, 150 mM NaCl, 10% Glycerol, 5 mM MgCl_2_, 1 mM DTT, 0.2% NP-40, protease inhibitor cocktail) was added to the ground tissue and vortexed. Centrifugation was performed at 18,000 g for 15 and 5 min subsequently to remove cell debris. Sample aliquots were either directly used for blue native-PAGE analysis, or immediately frozen in liquid nitrogen. Samples for SDS-PAGE were mixed with 3X SDS sample buffer (stock concentration 30% glycerol, 3% SDS, 93.75 mM Tris-Cl pH 6.8, 0.06% bromophenol blue) with 100 mM DTT at 70°C for 10 min.

### Protein extraction for large-scale immunoprecipitation

18 *N. benthamiana eds1* plants were infiltrated with Agrobacterium for transient expression. 2-3 leaves from each plant were infiltrated in 4-week old *N. benthamiana* plants. Leaves were harvested at 3 dpi, resulting in fresh weight of 15-20 g. These samples were immediately frozen in liquid nitrogen, and then ground with mortar and pestle, while maintaining the apparatus cold. Protein extraction buffer (50 mM HEPES pH 7.0, 150 mM NaCl, 10% Glycerol, 5 mM MgCl_2_, 1 mM DTT, 5 mM ascorbic acid, 0.2% NP-40, 0.2 mM PMSF, protease inhibitor cocktail, 2% PVPP) was added in 1 (tissue): 5 (buffer) ratio. Insoluble PVPP and cell debris were separated with centrifugation of 4,500 g for 15 min. After filtering the lysate with nylon mesh, the supernatants were centrifuged once more with 45,000 g for 45 min. The resulting supernatant was incubated with Flag M2 agarose beads (Merck A2220) for 1 h, and washed with extraction buffer without PVPP. Samples were eluted with 3xFlag peptides (Merck F4799) for 1 h.

### Blue native-PAGE

Protein extracts and immunoprecipitation (IP) eluates were added with 4× NativePAGE^TM^ Sample Buffer (Invitrogen^TM^, BN2003) and 1% Coomassie G-250 to a final concentration of 0.125%. Blue native-PAGE was performed with these samples as described in Ahn *et al*. (2023)^79^. NativeMark^TM^ Unstained Protein Standard (Invitrogen^TM^, LC0725) or SERVA Native Marker (SERVA, 39219.01) were used as molecular weight markers.

### Immunoblotting

Protein samples incubated with SDS sample buffer were loaded on SDS–PAGE gels and run at 90 V. After dye front reached the end, these gels were transferred with Trans-Blot® Turbo™ Transfer System (Biorad, #1704150) at conditions of 1.0 mA for 30 min onto PVDF membranes. Transferred membranes were blocked with 5% skimmed milk in TBS-T, and antibodies were added subsequently and incubated overnight at 4°C. The following antibodies were used; anti-Flag (Conjugated HRP, Merck, A8592), anti-Myc (Conjugated HRP, Merck, 16-213), anti-HA (Conjugated HRP, Roche, 12013819001), anti-StrepII (Merck, 71591-M), anti-Histone H3 (Agrisera, AS10 710), anti-PR1 (Agrisera, AS10 687), anti-Rabbit secondary (Conjugated HRP, Merck, AP132P). HRP Signals were detected using ECL substrates (Thermo Fisher, 34580). After detection, membranes were stained with Ponceau S solution (Merck, P7170) to use as loading control. PageRuler^TM^ Prestained Protein Ladder (Thermo Scientific, 26616, size range 10-180kDa) or Spectra™ Multicolor High Range Protein Ladder (Thermo Scientific, 26625, size range 40-300kDa) was used as molecular weight markers.

### Phylogenetic tree analysis of Arabidopsis TIR domains

The Arabidopsis Col-0 proteome (Araport11_pep_20220914) was downloaded from https://www.arabidopsis.org/. Interproscan v5^80^ was ran on each protein. Python and unix scripting was then used to select 140 proteins with hits to TIR domains (using terms “TIR” and “domain”). Proteins with more than one TIR domain in different regions were split based on having non-overlapping hits. TIRs for each proteins slice variants were manually inspected and only those non-overlapping sequences and unique sequences were kept. Highly truncated sequences were also removed. This left 152 unique TIR domains. ClustalW 2.1^81^ was used to create a 357 aa alignment. The alignment was trimmed of poorly aligned regions with trimAL v1.5^82^ to 108 aa. IQ-TREE v.2.3.3^83^ utilized to create a maximum-likelihood phylogenetic tree using model JTT+F+G4 with 1000 bootstraps. The structure of each TIR domain was predicted with AlphaFold2^84^. Alignments with the TIR domain from Roq1 (7jlx: S11-F176) were performed using frtmalign^85^. Amino acids in the corresponding structural positions to the Roq1 catalytic glutamic acid (E86), conserved BB-loop aspartic acid (D45), the BB-loop (F43-K53) and surrounding region (E37-E58) were extracted from alignments using unix scripting. These results were visualized using R studio with packages ape, ggplot2, ggtree, ggmsa and ggnewscale^86–90^. The tree was rooted to divergent TIR-N proteins AT5G56220.1 and AT4G23440.1. Code for these analysis are available at https://github.com/michhulin/Pseudomonas/tree/main/tirs.

### Sample Preparation for Mass Spectrometry

To identify the disulfide bonds in RRS1-R/RPS4 complex, 4-5 week-old N. benthamiana plants were used for infiltrating 3xFlag-tagged RPS4 and 4xMyc-tagged RRS1-R, sampled 3 dpi. Protein extraction was carried out with the following buffer: 50 mM HEPES pH 7.0, 150 mM NaCl, 10% Glycerol, 5 mM MgCl2, 10 mM NEM (N-ethylmaleimide), 0.2% NP-40, 0.2 mM PMSF, protease inhibitor cocktail, 2% PVPP. RPS4 were enriched using anti-Flag M2 beads (A2220, Merck), and eluted using 3xFlag peptide (F4799, Merck). Samples were incubated with SDS only and fractionated using SDS-PAGE. Gels that have corresponding size to RPS4 and RRS1-R were excised and in-gel digested by trypsin. Extracted peptides were measured for mass spectrometry analysis.

### LC-MS/MS Analysis

Extracted peptides were measured by Orbitrap Fusion in data-dependent acquisition method. The MS/MS fragmentation used both CID and HCD in parallel with each precursor ion selected for fragmentation. Raw files were peak-picked by MSConvert (Proteowizard project), searched by Mascot (Matrix Science) with added settings for disulfide crosslinks. The data could not be filtered to 1%FDR as current version of Mascot v2.8 does not support this validation. The mass spectrometry proteomics data have been deposited to the ProteomeXchange Consortium via the PRIDE^91^ partner repository with the dataset identifier PXD062781 and 10.6019/PXD062781.

### NADase activity assay

Fluorescence emitted from εNAD^+^ catalysis was measured to examine the NADase activity of the purified RRS1-R/RPS4 complex, as described in Tamulaitiene et al. (2024) ^92^. Briefly, RRS1-R/RPS4 complex was purified with anti-Flag M2 beads from transient expression in *N. benthamiana eds1* plants and eluted with 3xFlag peptide. 15 µl eluates were mixed with 100 μM εNAD^+^ (Merck, N2630) with or without 1 mM ATP. Reactions were measured using Spectra ID5 Max for in 1-min intervals for 1 h. 3 technical replicates were measured for each biological replication.

### Structural modelling

Structural models of the TIR domains were generated using Google Deepmind’s AlphaFold 3 server ^93^, using TIR domain sequences of RPS4 and RRS1-R. pTM and ipTM values were also taken from the AlphaFold 3 server. Structure alignments to determine if tetramers formed a ROQ1-like interface (PDB ID: 7JLX) and visualisation was done using UCSF ChimeraX ^94^. UCSF ChimeraX is developed by the Resource for Biocomputing, Visualization, and Informatics at the University of California, San Francisco, with support from National Institutes of Health R01-GM129325 and the Office of Cyber Infrastructure and Computational Biology, National Institute of Allergy and Infectious Diseases.

## Supplemental information

Document S1. Figures S1-S7

**Table S1.**

List of constructs generated in this study.

**Table S2.**

List of organisms and genetically modified organisms generated in this study.

**Table S3.**

List of Arabidopsis Col-0 TIR domains and their sequences.

**Table S4.**

List of RPS4^C783^-linked disulfide bonded peptides identified.

## Notes

### Competing Interest Statement

The authors have declared no competing interest.

