## Supplemental figures S1-S7 for "Recognition-dependent activation of the RRS1-R/RPS4 immune receptor complex"

#### This PDF file includes:

Figures S1-S7 and accompanied figure legends

### Figure S1

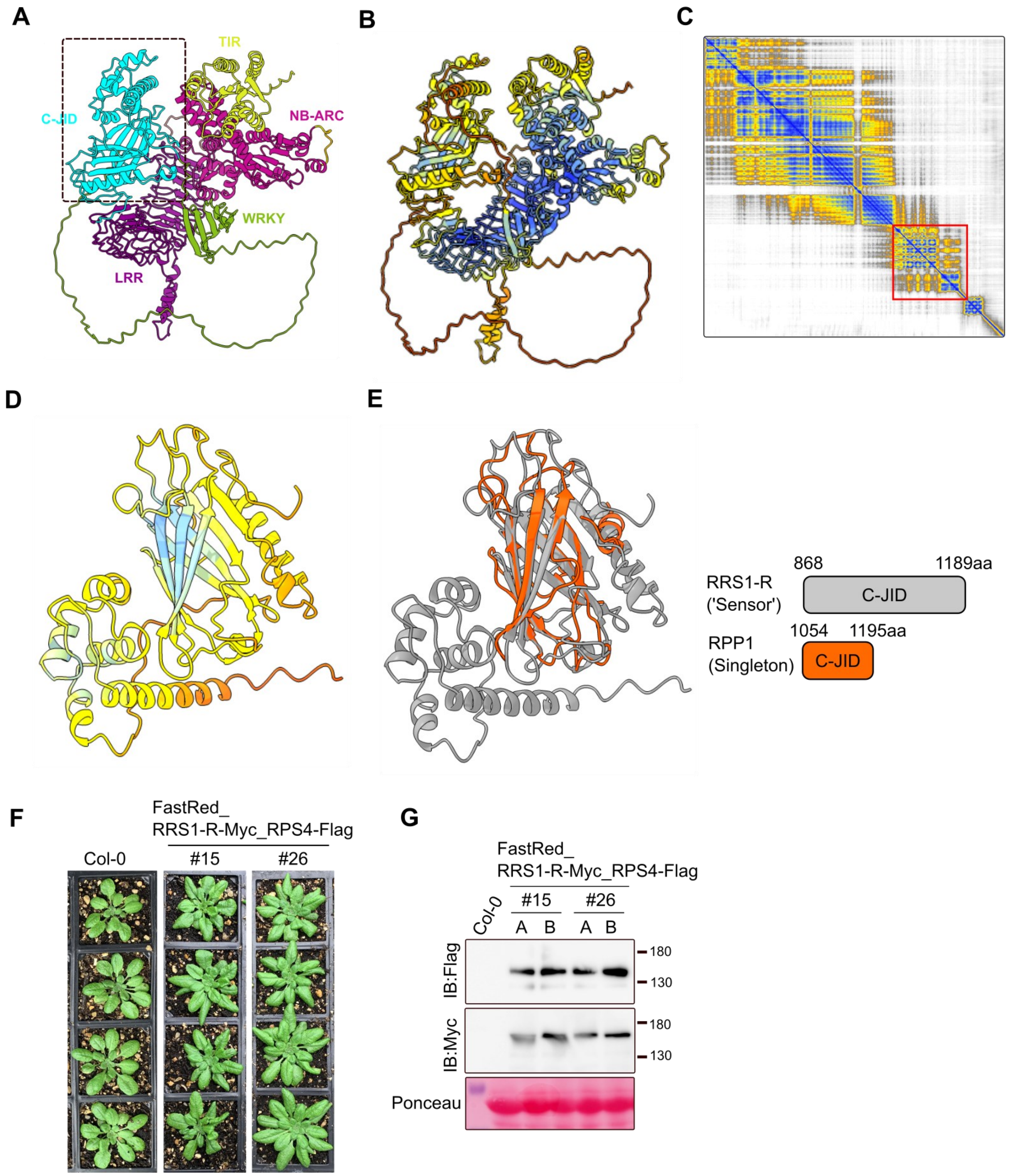


**Figure S1.**

A. RRS1-R Domain 4 is structurally predicted as C-JID. AlphaFold2 structure prediction of RRS1-R (EMBL AlphaFold Protein Structure Database; AF-C4B7M6-F1-v4) with domains in different colors for identification.

B, C. pLDDT scores for RRS1-R protein structure model from A. Red rectangle in C indicates the C-JID of RRS1-R.

D. Partial structure of RRS1-R C-JID (from A) colored with pLDDT scores.

E. RRS1-R C-JID (light grey) is superimposed with RPP1 C-JID (orange, PDB ID 7CRB) with Matchmaker function in ChimeraX. RMSD value is 274.599. Domain boundaries and length are shown on the right.(continued)

F. Growth phenotype of RRS1-R-Myc, RPS4-Flag overexpressing Arabidopsis transgenic line. 2 lines overexpressing RRS1-R and RPS4 were chosen and grown for 4 weeks. Col-0 plants sown together as control. Similar results were observed at least three independent times.

G. Protein extraction of RRS1-R-Myc, RPS4-Flag overexpressing Arabidopsis transgenic lines. Leaves from two independent plants of each transgenic line were collected from F, and proteins were extracted and loaded on SDS-PAGE. Protein extracted from Col-0 serve as negative control. Ponceau staining of the SDS-PAGE serve as loading control. Molecular weights of markers are shown on the right.

### Figure S2

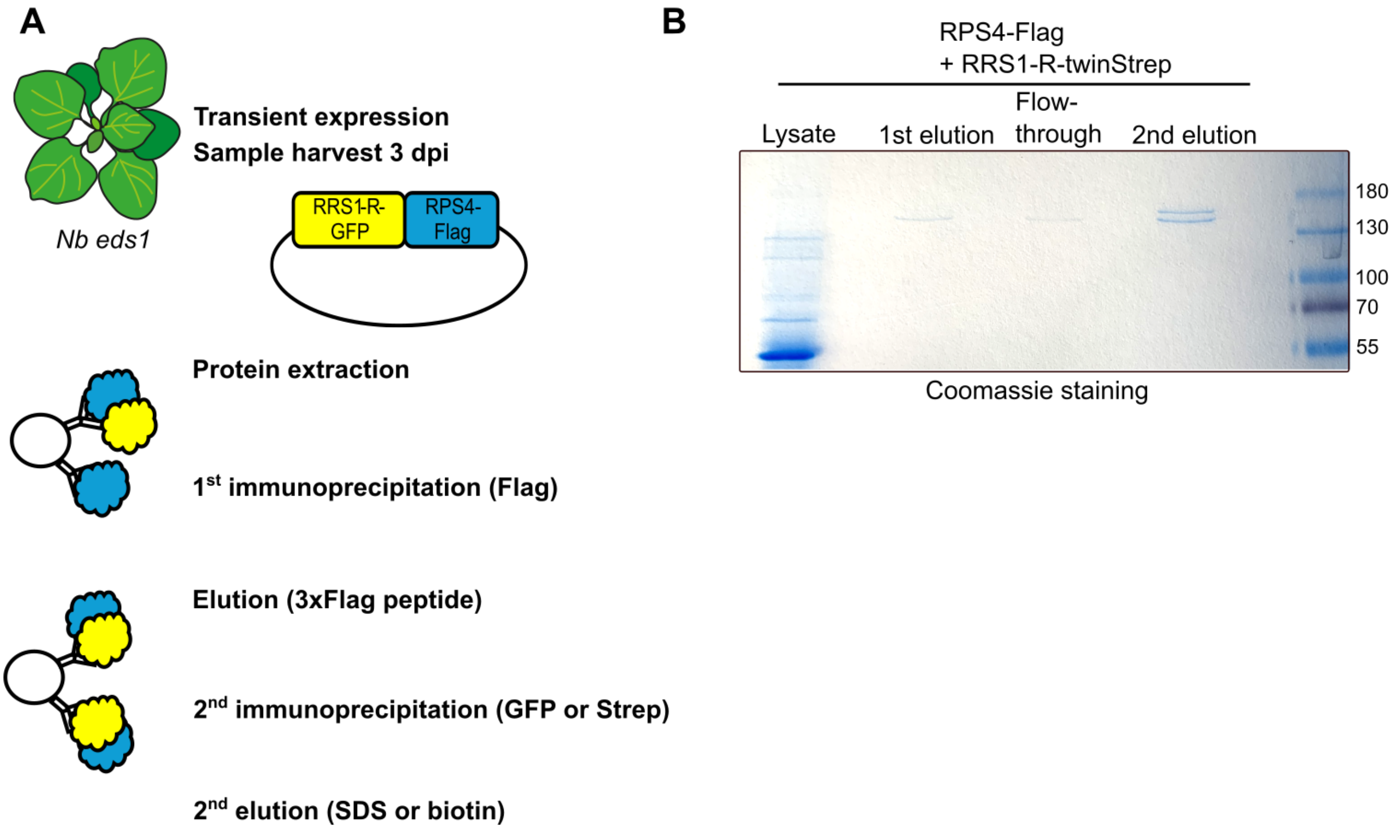


**Figure S2.**

A. Schematic diagram of tandem-affinity purification from N. benthamiana transient expression.

B. Tandem affinity purification improved purification yield of RRS1-R/RPS4 complex. Proteins were extracted from *N. benthamiana eds1* plants transiently expressing RPS4-Flag with RRS1-R-twinStrep. The protein extracts were immunoprecipitated using anti-Flag beads and eluted with 3xFlag peptide. The eluates were subsequently enriched with Strep-tactin resin, and bound protein complex were eluted with 10 mM biotin. Coomassie staining was performed with lysate, 1$^{st}$ elution product, supernatant after 2$^{nd}$ immunoprecipitation (flow-through), and the 2nd eluate product with biotin elution. Similar results were obtained in at least three biological replicates. Protein ladder is shown and size is indicated on the right.

#### Figure S3

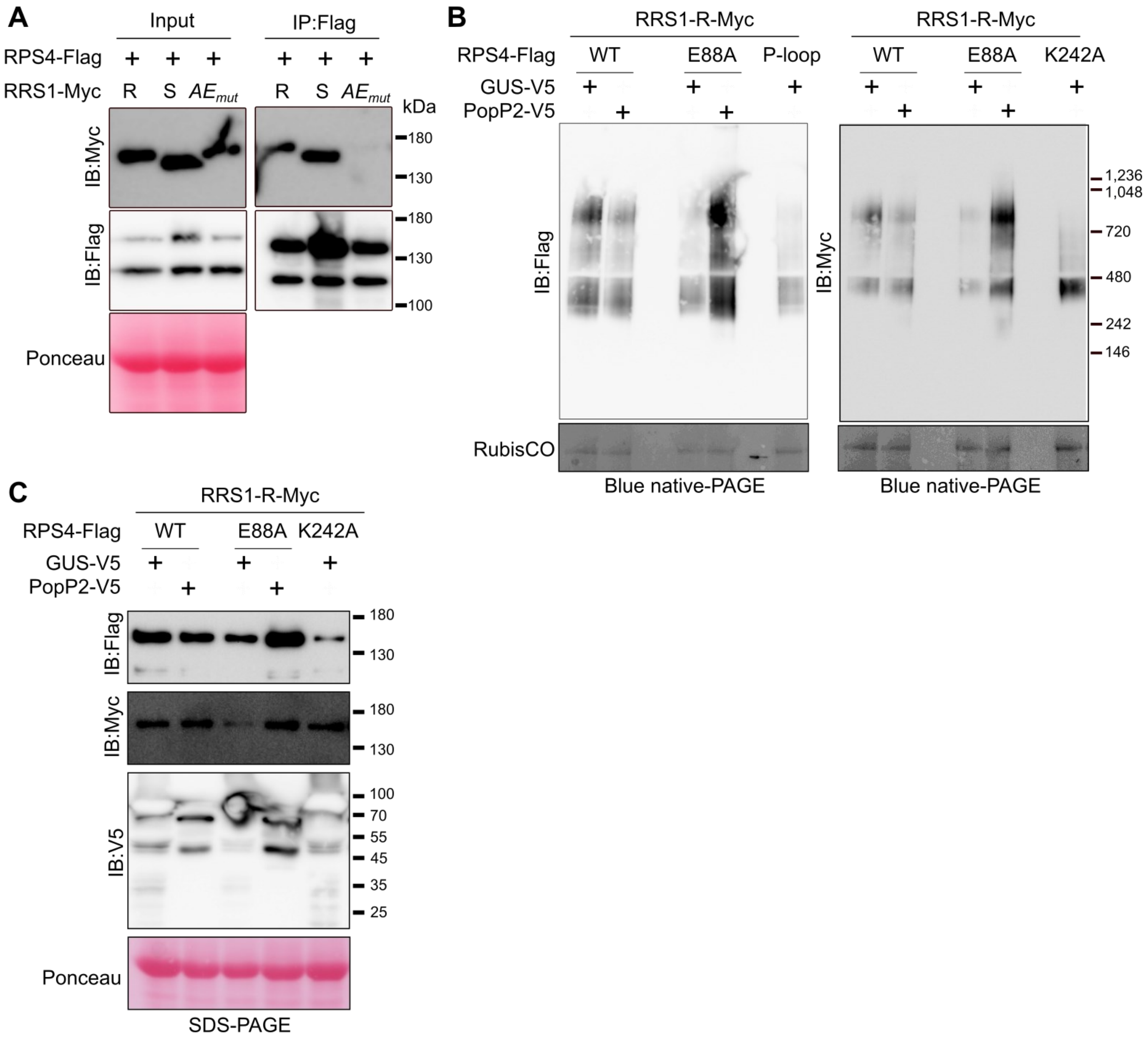


**Figure S3.**

A. RRS1-R-Myc AE interface mutant lost interaction with RPS4-Flag. Protein extracts from *N. benthamiana eds1* plants transiently expressing wild-type RPS4-Flag with wild-type or mutant RRS1-Myc were immunoprecipitated with anti-Flag beads. SDS-boiled input protein extract and IP eluate samples were loaded onto SDS-PAGE. Ponceau staining serve as loading control. Molecular weight markers are shown on the right. Experiments were performed at least three times with similar results.

B. RRS1-R/RPS4 complex does not alter stoichiometry upon activation upon co-expression of effector and loss of NADase activity. Proteins were extracted from *Nb eds1* plants expressing RPS4 wild-type or E88A mutant, with RRS1-R and in the presence or absence of effector PopP2. RPS4 P-loop mutant ($RPS4^{K242A}$) that is unable to form oligomer serve as negative control. Protein lysates were loaded on blue native-PAGE, followed by immunoblotting. Similar results were observed at three times, of which representative image is shown. Molecular weights of markers are shown on the right.

C. Protein samples extracted from B were incubated with 3XSDS sample buffer and 100 mM DTT at 70ºC, and loaded on SDS-PAGE. Ponceau staining serve as loading control. Molecular weight markers are shown on the right. Experiments were performed at least three times with similar results.

### Figure S4

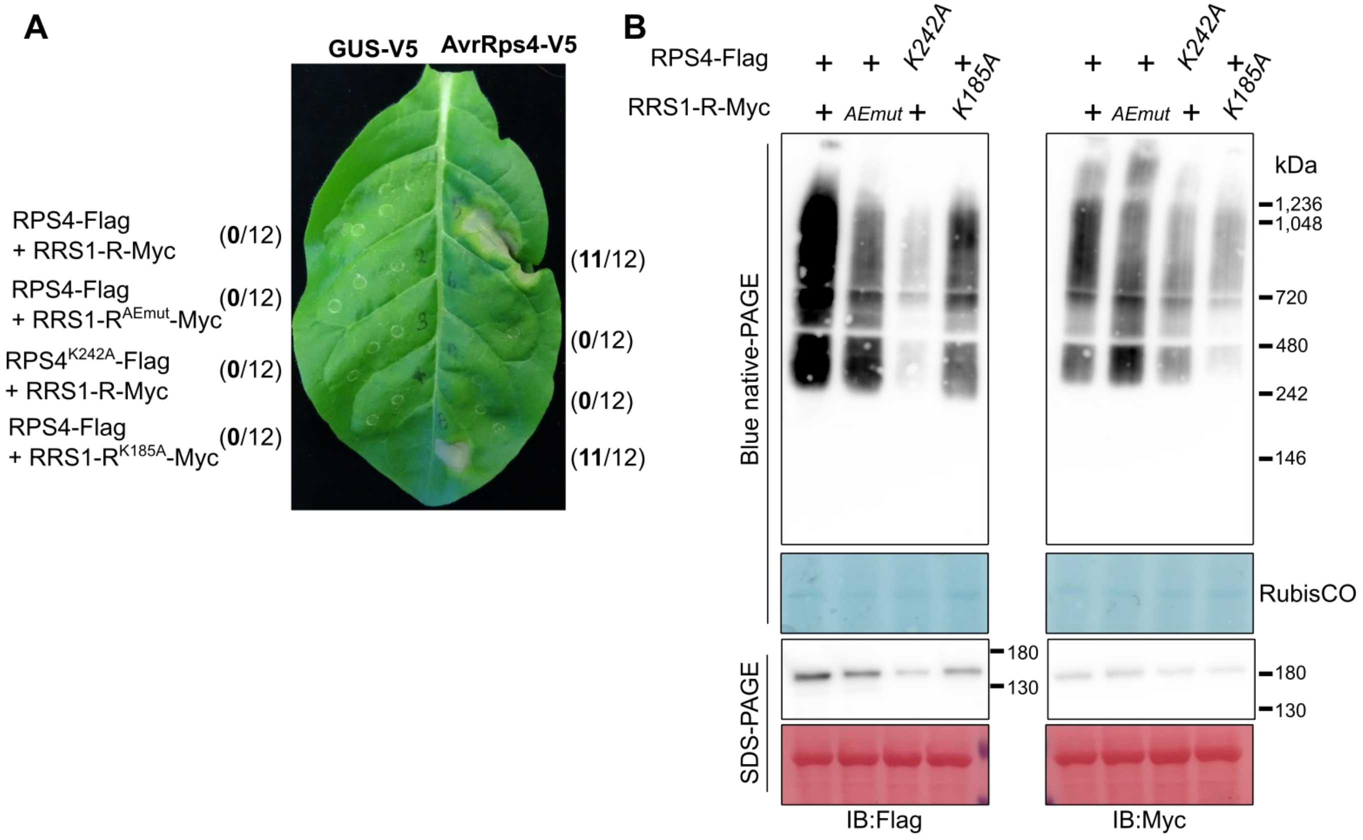


**Figure S4.**

A. Mutation of RPS4 P-loop (RPS4$^{K242A}$) leads to loss of effector-dependent HR by RRS1-R/RPS4. *N. tabacum* cv. *Petit Gerard* plants were infiltrated with RRS1-R wild-type or mutants, with RPS4 wild-type or RPS4 mutants in the presence or absence of effector AvrRps4, and tested for HR. Images were taken at 5 dpi, and at least 3 leaves were used for each replicate. Total number of cell death occurrences are indicated in parentheses.

B. P-loop mutation of RPS4 disrupt RRS1/RPS4 complex formation, but not P-loop mutation of RRS1. RPS4 and RRS1-R and its mutant derivatives transiently expressed in *Nicotiana benthamiana eds1* were loaded on BN-PAGE, followed by immunoblotting. Same samples were loaded on SDS-PAGE, followed by immunoblotting. RubisCO for BN-PAGE, and Ponceau staining for SDS-PAGE, serves as loading control. Remaining lysates were incubated with SDS and DTT and loaded on SDS-PAGE, followed by immunoblotting. Similar results were obtained at least three times, of which representative image is shown.

**Figure S5**

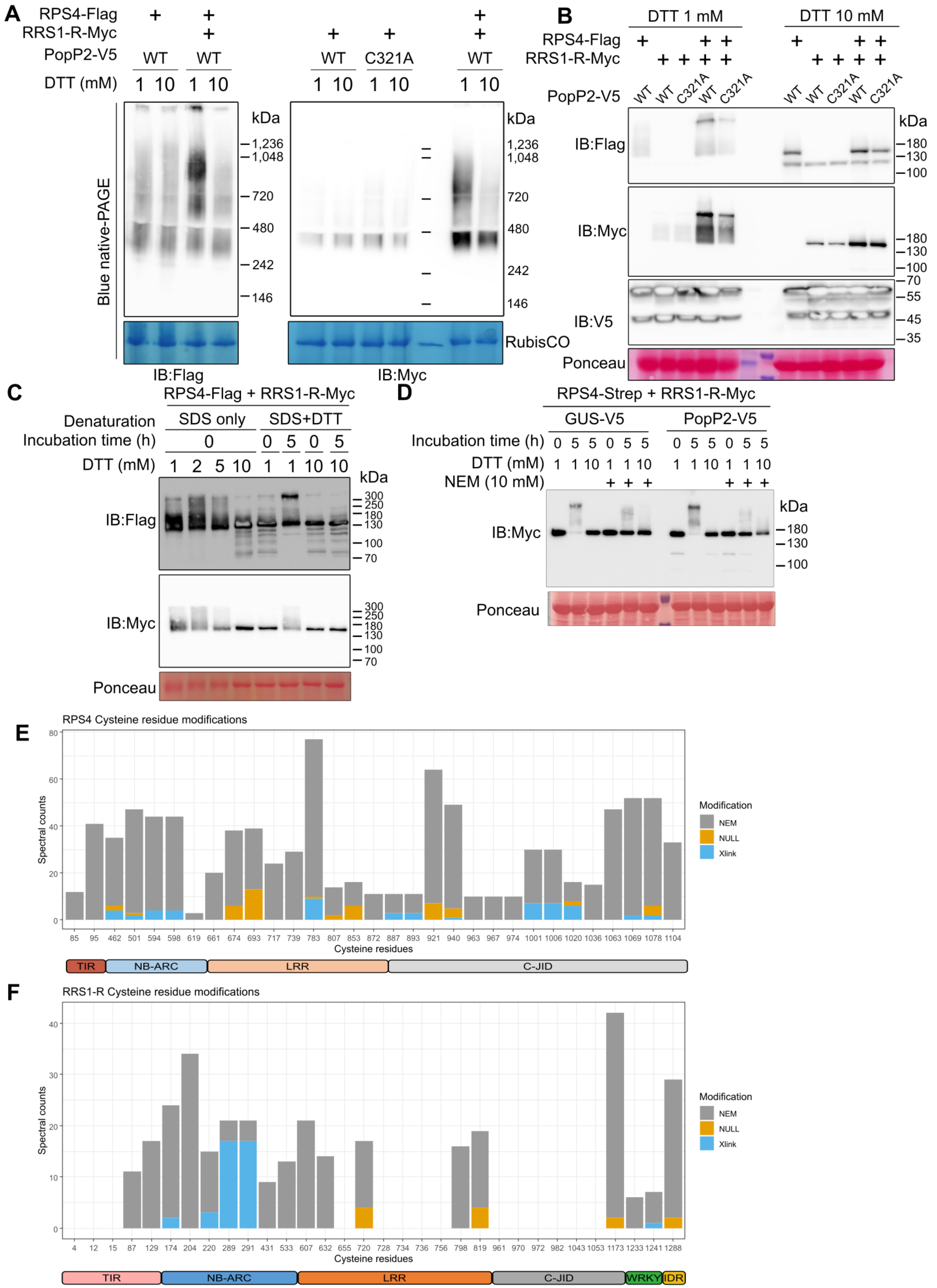
A
RPS4-Flag
RRS1-R-Myc
PopP2-V5
WT
C321A
DTT (mM)
1 10
Blue native-PAGE
kDa
1,236
1,048
720
480
242
146
RubisCO
IB:Flag
IB:Myc
B
DTT 1 mM
DTT 10 mM
RPS4-Flag
RRS1-R-Myc
PopP2-V5
IB:Flag
IB:Myc
IB:V5
Ponceau
180
130
100
70
55
45
35
C
RPS4-Flag + RRS1-R-Myc
Denaturation
SDS only
SDS+DTT
Incubation time (h)
DTT (mM)
300
250
IB:Flag
IB:Myc
Ponceau
D
RPS4-Strep + RRS1-R-Myc
GUS-V5
PopP2-V5
NEM (10 mM)
IB:Myc
Ponceau
E
RPS4 Cysteine residue modifications
Spectral counts
Cysteine residues
85 95 462 501 594 598 619 661 674 693 717 739 783 807 853 872 887 893 921 940 963 967 974 1001 1006 1020 1036 1063 1069 1078 1104
Modification
NEM
NULL
Xlink
TIR
NB-ARC
LRR
C-JID
F
RRS1-R Cysteine residue modifications
Spectral counts
Cysteine residues
4 12 15 87 129 174 204 220 289 291 431 533 607 632 655 720 728 734 736 756 798 819 961 970 972 982 1043 1053 1173 1233 1241 1288
Modification
NEM
NULL
Xlink
TIR
NB-ARC
LRR
C-JID
WRKY
IDR

**Figure S5.**

A. RRS1-R/RPS4 complex is dissociated under high reducing conditions. RRS1-R and RPS4 transiently expressed with PopP2 (wild-type or mutant C321A) in *N. benthamiana eds1* leaves were extracted with high (10 mM DTT) or low (1 mM DTT) reducing conditions, loaded on BN-PAGE, followed by immunoblotting. RubisCO complex observed on blue native-PAGE gel serves as loading control. Molecular weights of markers are shown on the right of each panel. Similar results were obtained at least three times, of which representative image is shown.

B. Protein extracts from A were incubated with SDS only, and loaded on SDS-PAGE, followed by immunoblotting. Ponceau staining of the SDS-PAGE serve as loading control. Molecular weights of markers are shown on the right. Similar results were obtained at least three times, of which representative image is shown.

C. Protein extracts from *N. benthamiana eds1* leaves transiently expressing RRS1-R-Myc and RPS4-Flag were extracted using various concentrations of DTT, and incubated on ice post-extraction. These samples were then denatured using SDS only or with SDS and DTT, and loaded on SDS-PAGE. Ponceau staining serve as loading control. Molecular weights of markers are shown on the right. Similar results were obtained at least three times, of which representative image is shown.

D. Protein extracts from *N. benthamiana eds1* leaves transiently expressing RRS1-R-Myc and RPS4-Strep with GUS (beta-glucuronidase)-V5 or PopP2-V5 were incubated on ice with varying reducing conditions. 10 mM NEM was added to the extract prevented formation of slower-migrating RRS1-R proteins under prolonged incubation conditions. Ponceau staining of the SDS-PAGE serve as loading control.

E. Graphical summary of disulfide bonds identified in RPS4 from LC-MS/MS. Peptide spectral counts with cysteine residues that are either modified with NEM (grey), crosslinked with another cysteine residue (Xlink; blue), or with no modification (Null; orange) were counted and plotted. Total number of spectral counts out of 2 individual biological replicates are shown. Position of each domains are shown in graphics below.

F. Graphical summary of disulfide bonds identified in RRS1-R from LC-MS/MS. Peptide spectral counts with cysteine residues that are either modified with NEM (grey), crosslinked with another cysteine residue (Xlink; blue), or with no modification (Null; orange) were counted and plotted. Total number of spectral counts out of 2 individual biological replicates are shown. Position of each domains are shown in graphics below. For cysteine residues with no counts, no peptides corresponding to the sequences were detected.

#### Figure S6

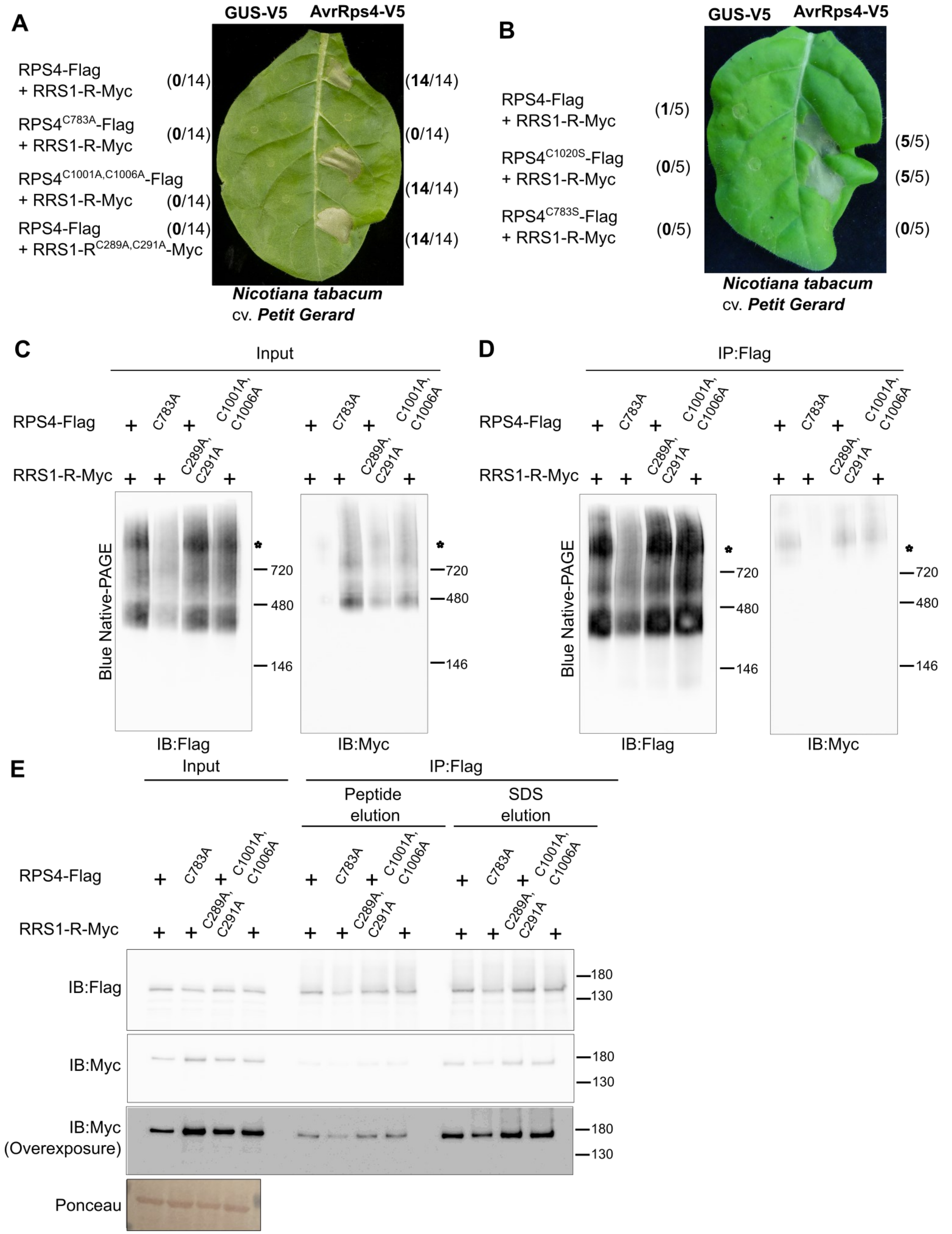


**Figure S6.**

A. RPS4 C783 mutation leads to loss of effector-dependent activation. Cysteine residues of RPS4 or RRS1-R identified to form disulfide bonds were mutated to alanine, and tested for effector-dependent activation of HR. After transient infiltration of wild-type of mutant RPS4-Flag and RRS1-R-Myc proteins with or without AvrRps4-V5 into *N. tabacum* cv. *Petit Gerard* leaves, images were taken 5 dpi. The number of leaves displaying cell death out of total number of leaves infiltrated are shown in parentheses. (continued)

B. RPS4 C1020 mutation does not lead to loss of effector-dependent activation. C783 and C1020 residues of RPS4 identified to form disulfide bonds were mutated to serine, and tested for effector-dependent activation of HR. After transient infiltration of wild-type of mutant RPS4-Flag and RRS1-R-Myc proteins with or without AvrRps4-V5 into *N. tabacum* cv. *Petit Gerard* leaves, images were taken 5 dpi. The number of leaves displaying cell death out of total number of leaves infiltrated are shown in parentheses.

C. RPS4 and RRS1-R and its mutant derivatives transiently expressed in *Nicotiana benthamiana eds1* were loaded on BN-PAGE, followed by immunoblotting. Molecular weights of markers are shown on the right of the panel. Similar results were obtained at least three times, of which representative image is shown.

D. Protein extracts from *N. benthamiana eds1* plants transiently expressing wild-type RPS4-Flag with wild-type or mutant RRS1-Myc were immunoprecipitated with anti-Flag beads. Proteins were eluted with 3xFlag peptide, and eluates were loaded on BN-PAGE, followed by immunoblotting.

E. SDS-boiled input protein extract and IP eluate samples from D and E were loaded onto SDS-PAGE. Ponceau staining serve as loading control. Molecular weight markers are shown on the right. Myc blots with higher contrast is shown below and labelled (overexposure). Experiments were performed three times with similar results.

**Figure S7**

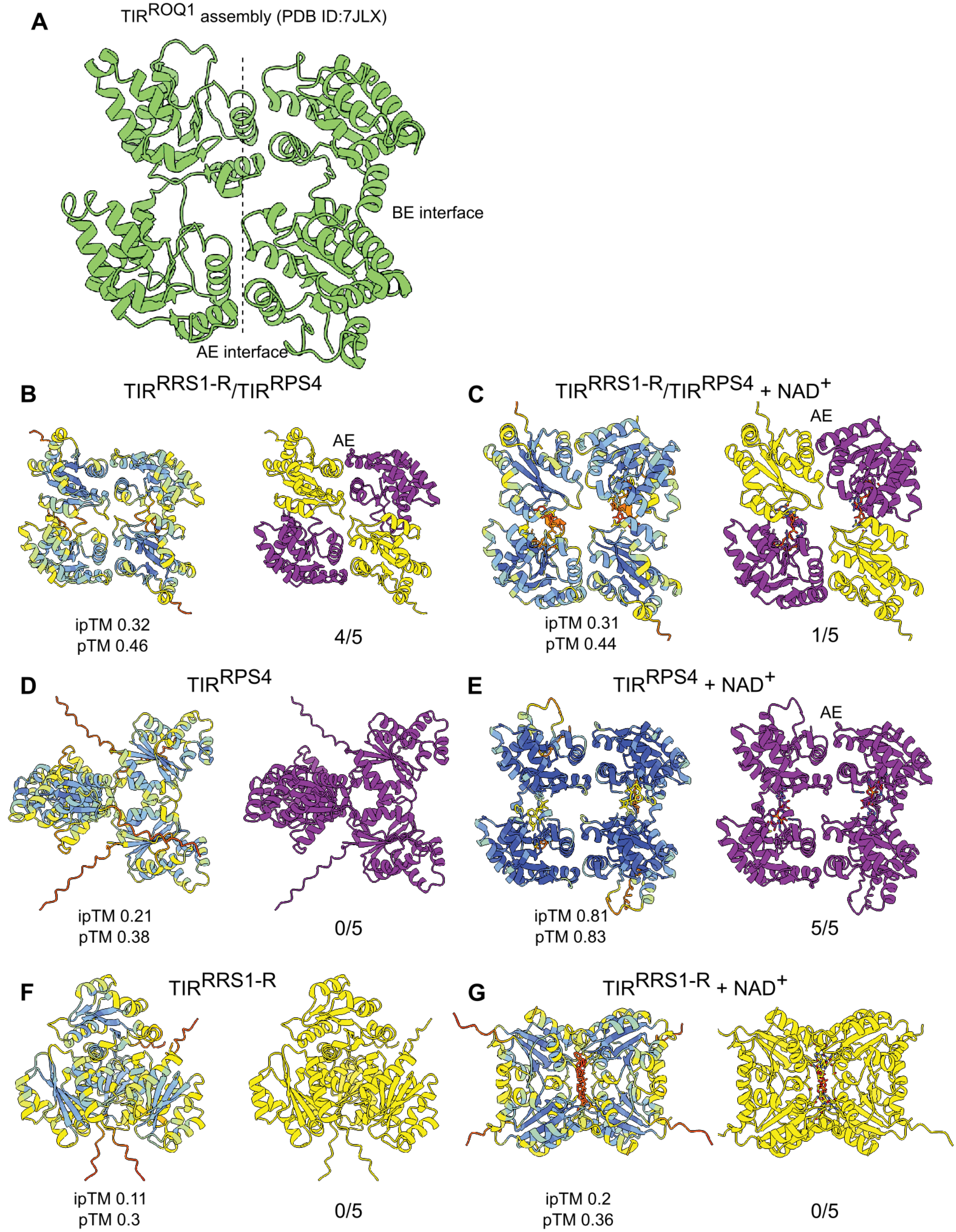


**Figure S7.** RRS1-R and RPS4 TIR domains can adopt a ROQ1 resistosome-like conformation both in the presence and absence of NAD+.

A. Cryo-EM structure of ROQ1 TIR domains (PDB ID: 7JLX). AE and BE interfaces labelled between monomers comprising the interface. (continued)

B-G. Alphafold3 models of $TIR^{RPS4}$ and $TIR^{RRS1}$ domains (B), $TIR^{RPS4}$ and $TIR^{RRS1}$ with NAD+ (C), $TIR^{RPS4}$ (D), $TIR^{RPS4}$ with NAD+ (E), $TIR^{RRS1}$ (F) and $TIR^{RRS1}$ with NAD+ (G). In each panel, the pLDDT colouring is shown on the left, with the ipTM (interface predicted temple modelling score) and pTM (predicted template modelling score) values shown below each model. On the right of each panel, $TIR^{RPS4}$ is coloured purple, and $TIR^{RRS1}$ coloured yellow, with the number of Alphafold3 models assuming a ROQ1 TIR-like assembly below each model. Figures prepared using ChimeraX, models, and pTM and ipTM scores, generated using Alphafold3 server.
